# Esca Disease triggers local transcriptomic response and systemic DNA methylation changes in grapevine

**DOI:** 10.1101/2025.08.11.669596

**Authors:** Margot MJ Berger, Virginie Garcia, Bernadette Rubio, Giovanni Bortolami, Gregory Gambetta, Chloé EL Delmas, Philippe Gallusci

## Abstract

Woody plants such as grapevine are vulnerable to trunk diseases caused by pathogens that colonize the wood, disrupt vascular function, and induce recurrent leaf symptoms associated with major metabolic disturbances and canopy decline. Over time, these diseases can irreversibly alter plant physiology and phenotype, ultimately reducing vine longevity. One of the most predominant of such diseases is esca, which is a major cause of vineyard dieback, with rising incidence worldwide over the past decade. However, the molecular mechanisms underlying esca symptom development remain unclear. In this study, we leveraged the heterogeneous expression of esca-symptoms within individual grapevines to investigate molecular responses in both symptomatic and asymptomatic leaf tissues. By combining metabolite profiling, RNA-seq and whole genome bisulfite sequencing, we show that metabolic alterations and extensive transcriptomic reprogramming are restricted to symptomatic leaves and are partially associated with local changes in DNA methylation. Asymptomatic leaves display distinct DNA methylation changes, some of which are shared with symptomatic tissues, suggesting a systemic response to the disease at the epigenetic level. Notably, subset of these methylation marks are observable prior to symptom emergence, highlighting the potential of epigenetic biomarkers for the early detection of trunk diseases in perennial plants.

## Introduction

Plants are sessile organisms that need to adapt to a constantly changing environment. As such they have developed a wide variety of mechanisms that drive their responses to stresses and adaptation to local environments (VanWallendael *et al*., 2019) among which epigenetic mechanisms play a central role (Mladenov *et al*., 2021). Originally defined by C Wadington as ‘‘the branch of biology which studies the causal interactions between genes and their products, which bring the phenotype into being” (Waddington, 2012), epigenetics now refers to the study of changes in genome functioning that are mitotically and/or meiotically heritable or stable and that do not entail a change in DNA sequence (Wu and Morris, 2001). Epigenetic modifications include histone post-translational modifications (HPTMs), DNA methylation, histone variants and require small RNAs (Zhang, 2008). DNA methylation is the most extensively characterized epigenetic mark to date (Zhang, Lang and Zhu, 2018). In plants, DNA methylation occurs at cytosines in the symmetrical CG, and CHG, and in the non-symmetrical CHH sequences contexts (where H is A, C, or T) (Zhang, Lang and Zhu, 2018). Methylation is maintained by context-specific DNA methyltransferase and can be removed by DNA demethylases or by passive dilution after DNA replication (Liu and Lang, 2020). De novo methylation is established by the RNA directed DNA Methylation (RdDM) pathway and small interfering RNAs (siRNAs) in all sequence contexts (Zhang, Lang and Zhu, 2018).

Recent works have highlighted the importance of epigenetic mechanisms in the response of plants to (a)biotic stresses (Fortes and Gallusci, 2017; Alonso, Ramos-Cruz and Becker, 2019; Joseph and Shah, 2022). To date, studies performed on model plants, mainly annuals such as *Arabidopsis* (Yu *et al*., 2013) rice (Akimoto *et al*., 2007), and tobacco (Wada *et al*., 2004), have shown a general increase in DNA methylation when plants are under pathogen attack. Concomitantly, DNA demethylation occuring at promoters of genes involved in plant defense, has been associated with their expression (reviewed in (Hewezi *et al*., 2018; Joseph and Shah, 2022). In contrast, general epigenetic responses are less studied in woody perennials than in annuals, and this is especially true for epigenetic responses to pathogen infection. So far, studies, performed with Poplar and Picea have shown that epigenetic processes are involved in perennial plant development and their responses to environmental stresses (Yakovlev *et al*., 2012; Bräutigam *et al*., 2013), reviewed in (Albaladejo *et al*., 2019)). At the same time, woody perennials present specificities that cannot be addressed in annual plants. This includes the ability to remember the environmental conditions they have been facing over years (trans-annual memory, (Gallusci *et al*., 2023)), plant aging (Yao, Schmitz and Johannes, 2021). This is especially relevant when addressing diseases of perennial organs such as the trunk, and these diseases represent a major threat in the context of climate change for both wild and cultivated trees, including grapevine (Claverie *et al*., 2020; Martino, Spadaro and Guarnaccia, 2025).

In this context, grapevine is one of the most economically valuable fruit crops worldwide (Alston and Sambucci, 2019) and is facing several threats due to climate change and agricultural constraints. Since the 2000s, important yield losses and mortality increases have been observed in old-growth vineyards mainly in Europe and Mediterranean vine-growing regions. This decline has been associated with the increased incidence of grapevine trunk diseases, particularly esca, which have risen significantly over the past decade in vineyards across Europe, America, and South Africa, although other contributing factors may also be involved (Cloete *et al*., 2015; Mondello, Larignon, *et al*., 2018; Guerin-Dubrana, Fontaine and Mugnai, 2019; Claverie *et al*., 2020; Etienne *et al*., 2024). Esca is a complex disease caused by a consortium of pathogenic fungi from both ascomycete and basidiomycete families (Gramaje, Úrbez-Torres and Sosnowski, 2018). It is characterized by internal trunk necrosis (Maher *et al*., 2012), sometimes accompanied by foliar symptoms characterized by a typical “tiger-stripe” leaf scorch pattern observed in summer (Lecomte *et al*., 2024), as well as berry damage that can lead to significant yield losses (Mugnai, Graniti and Surico, 1999; Lecomte *et al*., 2012; Dewasme *et al*., 2022), both of which which are typically observed in plants older vines (i.e.> 7 years of age; (Mondello, Songy, *et al*., 2018). Interestingly, the fungi associated with esca disease which are located in the trunk, are not detected in current year shoots nor in symptomatic leaves (Bortolami *et al*., 2019; Gastou *et al*., 2025). In contrast, esca leaf symptoms have been associated with hydraulic failure caused by tyloses occluding the xylem vessels in current year shoots (Bortolami *et al*., 2023) and with a decrease in plant transpiration, leaf chlorophyll, and non-structural carbohydrate content (Bortolami *et al*., 2021). In addition, major metabolic changes occur in leaves and fruits of symptomatic plants as compared to non-symptomatic ones (Weiller *et al*., 2024).

Despite the precise knowledge of the esca disease phenotype, there is an extremely limited understanding of the molecular mechanisms underlying trunk disease symptom expression in grapevine, including esca. Molecular studies on esca have focused primarily on gene expression patterns associated with symptom development, particularly involving genes related to primary metabolism and physiological processes such as photosynthesis (Magnin-Robert *et al*., 2016; Ouadi *et al*., 2021). More recently, transcriptomic reprogramming was described in wood tissues inoculated with esca-associated pathogens under controlled conditions (Romeo-Oliván *et al*., 2024). Metatranscriptomic approaches have also been applied to characterize microbial mRNA populations in trunk tissues from field-grown vines with or without foliar symptoms (Morales-Cruz *et al*., 2018; Nerva *et al*., 2022). While a recent field-based transcriptomic study has explored grapevine responses to a different trunk disease (Patanita *et al*., 2025), no integrated molecular analysis has yet been conducted on esca-affected grapevines under vineyard conditions. As a result, there is still a lack of comprehensive understanding of the plant’s molecular responses to esca in its natural context, especially at the whole-plant and tissue-specific levels.

In this study, we analyzed grapevine that have consistently expressed esca-disease symptoms over the past five years (esca-plants). Leveraging the unique symptomatology of esca, which often results in both symptomatic and asymptomatic sectors within the same canopy, we developed an integrative approach that extends from the analysis of epigenetic landscapes, transcriptomic and metabolic profiles to plant phenotypes. Our findings revealed that while only symptomatic leaves (ES) presented significant changes in metabolite composition and extensive transcriptional reprogramming, alteration in the DNA methylation landscape occurred in all canopy sectors, irrespective of the presence of foliar symptom. Importantly, some of the esca associated epigenetic changes were also found in esca-plants before leaf symptoms were expressed, consistent with an epigenetic signature of esca-plants. As a conclusion, esca disease leads to an epigenetic remodeling of plants independently of the development of symptoms, which may provide an innovative strategy to identify plants that are infected by esca before any symptom is visible.

## Material and methods

### Plant material

Plants (*V.vinifera* L. cv Sauvignon blanc grafted on Fercal rootstock) were cultivated in Château Couhins (Villenave d’Ornon, France, 44°45’16.272”, - 0° 33’ 32.148”) since 2001, with a yearly monitoring of esca leaf symptom since 2012. Leaves were sampled in 2019 from three plants that have not shown any esca symptoms since 2012 (Control-plants), and three symptomatic plants (Esca-plants) that had expressed symptoms during three to four years between 2012 and 2019 (Table S1, Figure 1A). Each “Control” sample corresponds to 4 pooled leaves sampled from one single plant. For each Esca-plants, two different samples were collected: (1) a pool of 4 symptomatic leaves with typical tiger-striped features associated with esca expression (ES leaves), (2) a pool of 4 green non symptomatic leaves coming from asymptomatic canes of the same plant (EA leaves) (Figure 1A). Leaves were directly frozen in liquid nitrogen after leaf veins and petiole excision, and the removal of necrotic tissues in the case of ES leaves. Tissues were frozen-ground to a fine powder and stored at -80°C until use.

**Figure 1.**
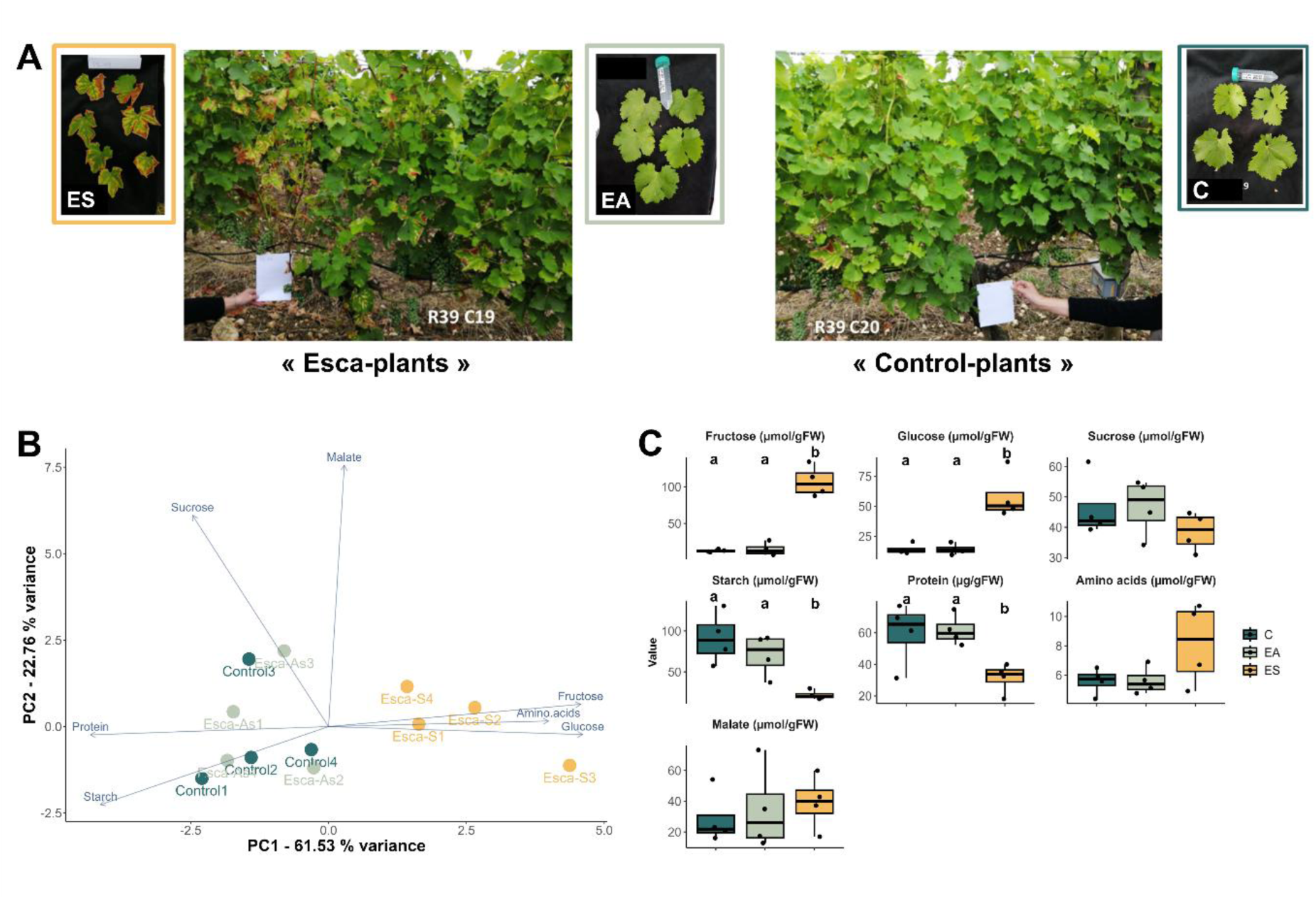
Phenotypes of plants grown in Vineyards: (**A**) Symptomatic (inset ES) and Asymptomatic (inset EA) leaves “Esca-plants” (plants that have expressed symptoms during 3 to 4 years between 2012 and 2019), and Control (inset C) leaves from “Control plants” (Plants that not expressed any symptoms during the same period; see methods). (**B**) Principal Component Analysis of ES, EA and C leaves based on the concentration of the seven metabolites measured in this study. (**C**) Concentration of primary metabolites in C, EA and ES leaves. C: Control leaf, ES: esca-plant symptomatic leaf, EA: esca-plant asymptomatic leaf

The plants used in greenhouse experiment were described in Bortolami et al., (2021) and correspond to ‘well-watered’ plants of *V.vinifera* L. cultivar Sauvignon blanc grafted on 101-14 Millardet et de Grasset (101-14 MGt) Nouvelle-Aquitaine Bordeaux (44°47′24.8″N, 0°34′35.1″W). Plants were monitored for esca symptoms in the vineyard from 2012 to 2017) before being transplanted into pots in 2018 and further monitored in 2018 and 2019 in greenhouse (Table S1). Potted plants without esca-symptoms monitored from 2012 to 2019 were named “Control-plants” (pC), while plants that expressed esca-symptoms (Table S1) were referred to as “Esca-plants” (pE). One young leave (4^th^ from the apex) per plant were sampled from 3 pC and 3 pE, at the beginning of June 2018 and 2019 before foliar symptoms expression (Figure S1), following the same procedure as for vineyard experiments.

### Nucleic acid extraction

RNA and/or DNA were extracted from approximately 200 mg (FW) leaf powder following the procedure described in (Berger *et al*., 2022).

### Targeted metabolites extraction and analysis

Metabolite extraction was performed in quadruplicate for each condition from frozen leaf powder. Extraction was performed as described in (Luna *et al*., 2020) at the *Bordeaux Metabolome Facility* (https://metabolome.u-bordeaux.fr/en/, Villenave d’Ornon, France) with the following modifications: (1) 20 mg of fresh frozen-powder (FW) of each sample were used for extraction, (2) ∼10 mg of polyvinylpolypyrrolidone (PVPP) were added in each tube prior to extraction to precipitate polyphenols. Analyses of soluble sugars (glucose, fructose, sucrose), starch, malate, citrate, total soluble proteins, total amino acids were conducted at *Bordeaux Metabolome Facility*. Measurements were based on couple enzyme assays as described previously in (Biais *et al*., 2014), except for total soluble proteins measured by Bradford assay (Bradford, 1976).

### RNA-sequencing

High throughput sequencing of RNA samples was performed using DNBSEQ-sequencing technology (pair-end, 150bp) service, provided by BGI-Genomics platform (http://bgi.com/). We generated 9 cDNA libraries corresponding to the three biological replicates for each condition of the vineyard experiment. Sequence alignment files were processed with the nf-core (nfcore-Nextflow-v23.10.0 ; (Ewels *et al*., 2020)) pipeline ‘rnaseq’ (3.14.0) for quality control (QC), trimming, alignment (STAR-RSEM route ; (Dobin *et al*., 2013)) (Table S2) and count matrix generation using *Vitis vinifera* reference genome PN40024 (PN40X, v4; (Velt *et al*., 2023). Differential expression analysis was performed using R (v 4.3.1) DESeq2 package (v 1.42.1), and Wald test (Love, Huber and Anders, 2014). We compared Transcriptomic profile of Esca-symptomatic (ES), Esca-asymptomatic (EA) and control (C) leaves as detailed in Figure 2A. As both ES and EA leaves came from the same plants, comparison ES-EA was performed using a multi-factor design aimed at estimating the effect of esca leaf symptom, while accounting for the paired-nature of the samples. Differentially expressed genes were identified using a False Discovery Rate (FDR) adjusted p-value threshold <0.05 and an absolute fold change >2 (l2fc >|1|). Functional description for each DEG was retrieved from ‘PN40024.v4’ reference genome (data provided by GRAPEDIA consortium, https://grapedia.org/).

**Figure 2.**
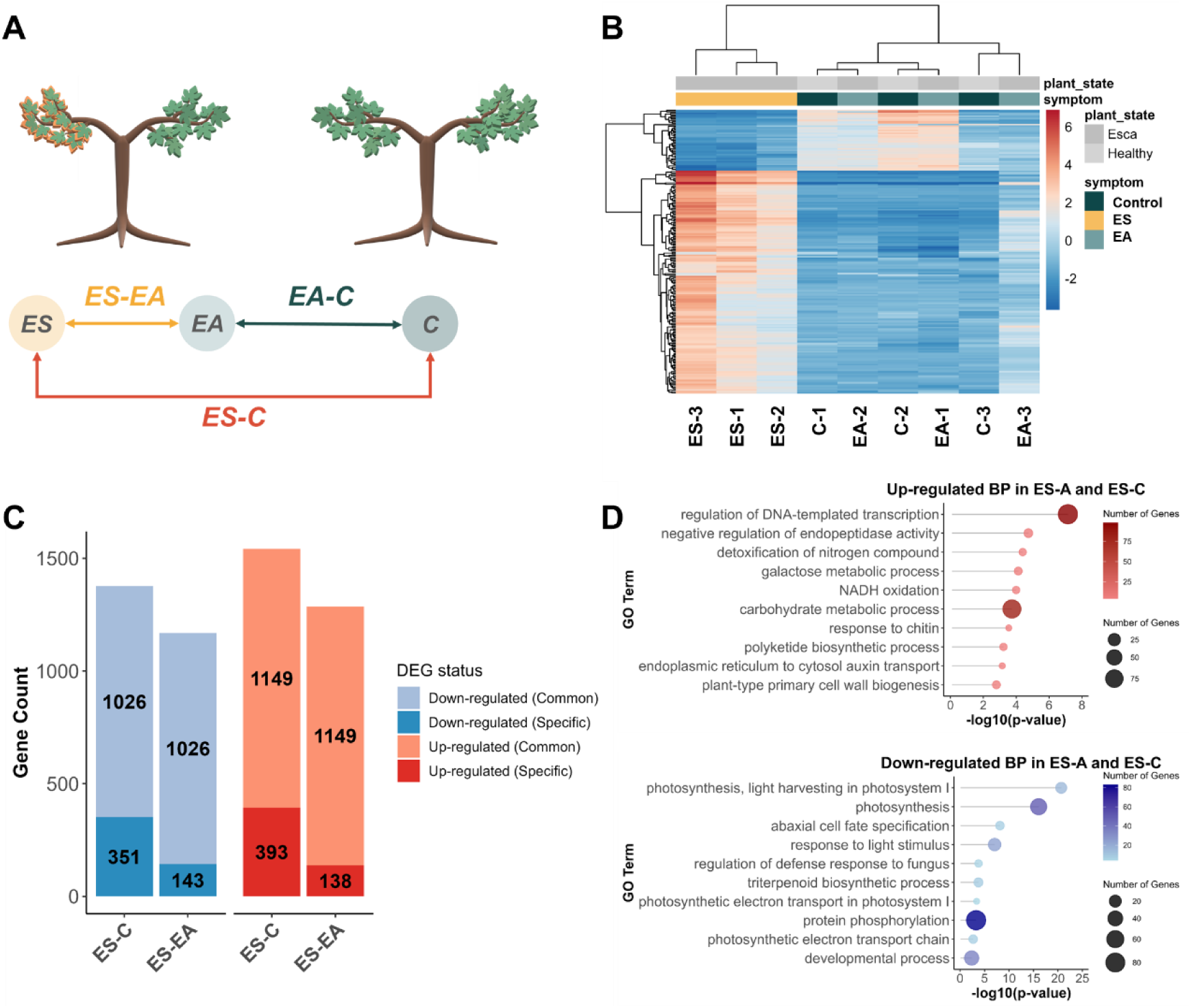
(**A**) Differential analyses were performed by comparing ES to EA leaves (ES-EA), ES to C leaves (ES-C), and EA to C leaves (EA-C) Analysis of the mRNA populations in EA, ES and C leaves. (**B**) Hierarchical clustering of EA, EA and C leaf samples based on the expression profile of the 200 genes with the highest difference in expression level between samples. (**C**) DEGs identified in ES-C and ES-EA comparisons. Up-regulated genes are indicated in red, and down-regulated ones in blue. Common DEGs are shown in light colors. (**D**) Gene Ontology analysis of up- and down-regulated DEGs common to the ES-EA and ES-C comparisons. C: Control leaf, ES: esca-plant symptomatic leaf, EA: esca-plant asymptomatic leaf

### Whole genome bisulfite sequencing

Whole genome bisulfite sequencing of DNA samples was performed with the DNBSEQ-sequencing technology (pair-end, 100bp, BGI-Genomics) with a mean sequencing depth ranging between 19.8 and 21.4X depending on samples (Table S3). Bisulfite conversion rate was calculated with Bismark tool (>99.1%) using the *Vitis vinifera* chloroplast complete genome (NCBI Reference Sequence NC_007957.1) which is un-methylated. Reads obtained for vineyard and greenhouse samples (14 Gb per sample) were further processed with nf-core (nfcore-Nextflow-v23.10.0)/Methylseq (v2.6.0) pipeline providing adapter sequence trimming, read alignment (Bismark v0.24.0, default parameters, mismatches set to 5) on the grapevine reference genome (PN40024, PN40X, Velt et al., 2023), and extensive quality control (QC). Only uniquely mapped reads were keep for further analysis. Global samples methylation levels at genome scale or specific genomic features (genes, TE) were represented with viewBS (v0.1.11). Before any further analysis, data were filtered to ensure that cytosine positions considered for analysis were present in every dataset with a minimum ‘coverage’ ≥ 5. Differential methylation analyses were performed using DSS (Dispersion Shrinkage for Sequencing data – v2.50.1) (Feng and Wu, 2019) to identify differentially methylated regions (DMRs) based on a Wald-test procedure. DMRs identification between non-paired samples (ES-C, EA-C, pE-pC, Figure 2A, Figure S1)) were performed with default parameters with subsequent steps of statistical analysis at each C locus by calling ‘DMLtest’ function (smoothing=TRUE), and DMR identification using ‘callDMR’ function (delta=0.1, p.threshold=0.01). For paired sample differential analysis (ES-EA, Figure 2A), we used DSS fixed effect model fitting ‘DMLfit.multiFactor’ with a ‘design’ accounting for leaf phenotype (ES or EA) and plant (Esca-plant 1, Esca-plant 2 or Esca-plant 3) from which both ‘ES’ and ‘EA’ leaves were sampled (formula=∼phenotype+plant). Statistical analysis at each cytosine position was subsequently performed with the function ‘DML.test.multifactor’ to identify differential methylation related to esca-symptom expression (term=”phenotype”), followed by DMRs identification using ‘callDMR’. Since ‘DML.test.multifactor’ function does not return relative methylation levels, we estimated relative methylation differences for each DMRs identified by (1) extracting methylated counts and total counts for every cytosine within the DMR for each sample, (2) calculating mean methylation by dividing ‘total methylated counts’ by the ’total counts’ values (3) estimating methylation difference by mean methylation level subtraction between conditions. To test whether methylation levels between comparison groups were statistically different, p-value for each DMRs were calculated using a t.test (paired=TRUE) considering methylation levels across the three biological replicates in each condition. The p-value was then adjusted using Benjamin-Hochberg method. DMRs showing 10% methylation difference associated with a p.adj ≤ 0.05 were considered for further analysis.

### Identification of potential epigenetic markers of esca

Two different approach were used to identify potential epigenetic markers of esca: (1) a ‘context-restrictive’ approach, by identifying context-specific (CG, CHG and CHH) overlapping positions of DMRs identified in the ES-C, EA-C, pE-pC 2018 and pE-pC 2019 comparisons (Figure 2A, Figure S1). (2) A ‘DMR-enrichment’ approach, by merging overlapping or nearby (distance < 500bp) DMRs identified in all comparisons (ES-EA, ES-C, EA-C, pE-pC 2018, pE-pC 2019 ; Figure 2A, Figure S1). Genomic regions presenting DMR accumulation (more than 2), irrespective of their methylation context, were considered as ‘enriched’. Overlapping and analysis were performed using the GenomicRanges (v.1.54.1) R (v4.3.1) package.

## Results

### Leaves with esca symptoms present specific gene expression profiles associated with metabolic changes

To evaluate the metabolic effects of esca on grapevine leaves, metabolite profiles were analyzed in leaves of esca-plants with (ES) or without (EA) symptoms and of control-plants (C) (Figure 1A). Principal Component Analysis (PCA) of the abundance of seven primary metabolites separate ES samples from all others (Figure 1B). The PC1, which account for 61.53% of the variance, is primarily influenced by fructose, amino acids, glucose, protein and starch levels. The ES leaves were characterized by an accumulation of glucose, fructose, and amino acids, while sucrose, starch and total protein levels decreased (Figure 1C). Metabolic profiles of EA and C leaves were similar, indicating that the metabolic changes observed in ES leaves were symptom-driven (Figure 1B, C). The RNAseq analyses were also performed to compare mRNA populations of ES, EA and C samples (Figure 2A, S2). Principal Component Analysis (PCA) of the transcriptomic profiles based on all expressed genes (Figure S2), and hierarchical clustering using the 200 genes presenting the highest variations in expression levels between samples (Figure 2B) showed that ES leaves were grouped and clearly separated from all other types of leaf. EA and C leaves have similar gene expression patterns with only one differentially expressed gene (DEG) (*Vitvi05g01942*, Early-nodulin-75-like, L2FC: -2.5, padj: 0.02).

By contrast, the mRNA population of ES leaves is clearly distinct from those of EA and C leaves with 2,456 and 2,919 DEGs respectively, equally distributed between up and down-regulated genes (Figure 2C). The majority of DEGs (∼ 75%) were common between the ES-EA and ES-C comparisons (Figure 2C), and present the same differential expression profiles. Gene Ontology (GO) performed on these common DEGs reveals that up-regulated genes in ES leaves are primarily enriched in biological processes related to the reprogramming of transcriptomic activity, biotic stress response, detoxification, and to the activation of stress-related metabolic pathways, while down-regulated genes were primarily enriched in biological processes related to photosynthesis, light harvesting, protein phosphorylation, development, and hormone synthesis (Figure 2D). Only 281 and 743 DEGs are specific to the ES-EA and ES-C comparisons respectively. DEGs specific to the ES-EA comparison are enriched in carbon metabolic processes, and those specific to the ES-C comparison in transcription, auxin regulation, and signaling (Figure S3), demonstrating specific transcriptomic profiles associated with the plant state from which asymptomatic leaves were sampled.

In summary, symptomatic leaves were characterized by specific metabolic and transcriptomic reprogramming, whereas asymptomatic leaves displayed comparable profiles regardless of the plant of origin.

### Leaves of plants expressing Esca symptoms exhibit DNA methylation changes indicative of both localized and systemic responses

Whole-genome bisulfite sequencing revealed similar global methylation levels and patterns across ES, EA, and control leaves, with slight, non-significant differences in CG, CHG and CHH methylation (see Methods, Figure S4, Table S3). PCA showed limited overall variation, but indicated a distinction in clustering patterns across methylation contexts, with ES samples more dispersed in CG, and esca-plant samples (ES and EA) clustering together in CHG and CHH (Figure S5).

We identified differentially methylated regions (DMRs) to assess the impact of Esca symptoms on leaf methylation profiles. In total 669 DMRs were found in ES-C, 563 in ES-EA and 303 in EA-C comparisons, indicating that symptomatic leaves (ES) differ substantially from asymptomatic ones (EA and C), regardless of plant states (Figure 3A).

**Figure 3.**
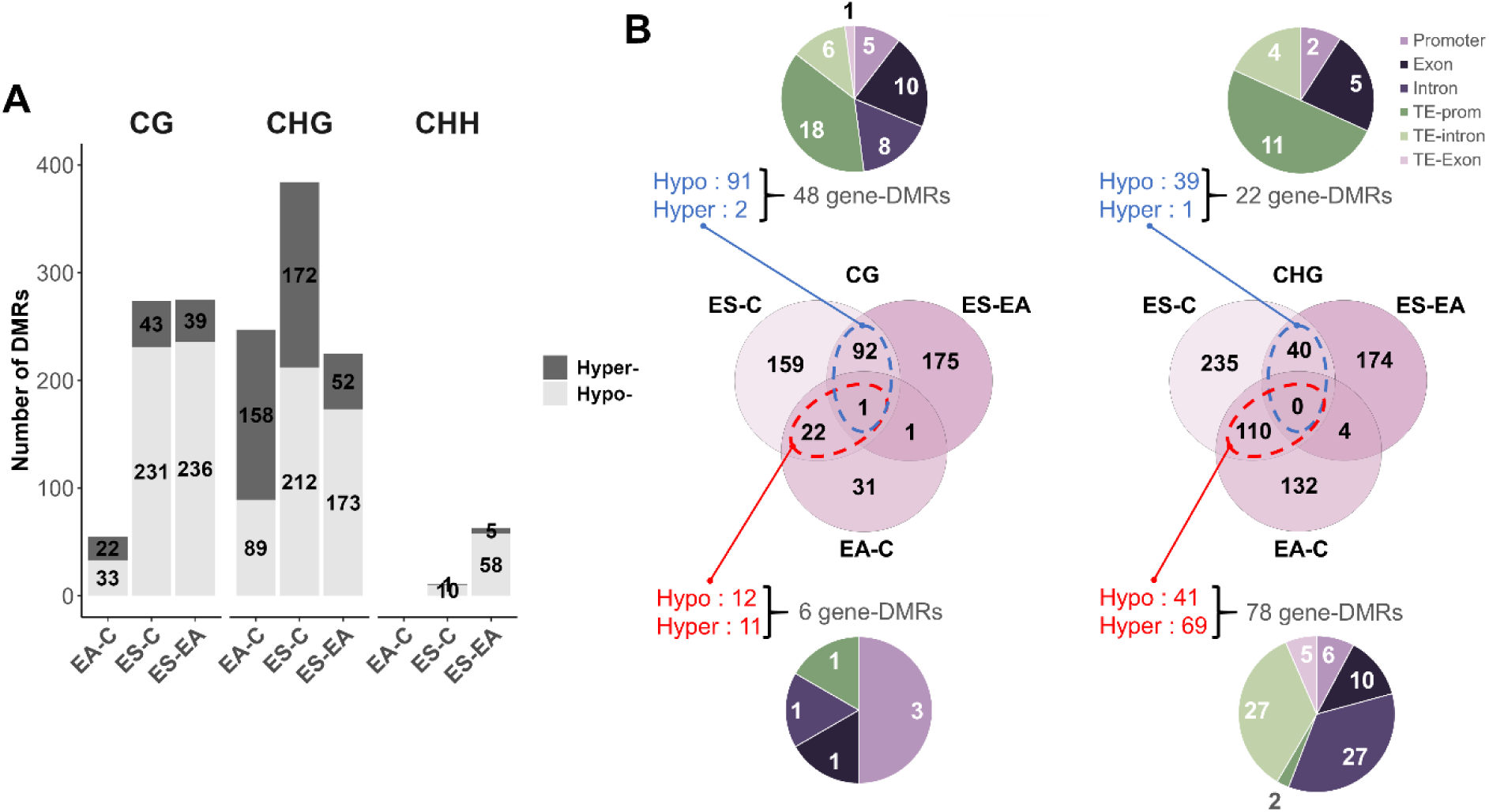
Changes in DNA methylation identified in ES, C and EA in leaf samples. (**A**) Number and methylation status of differentially methylated regions (DMRs) identified in ES-C, EA-C and ES-C comparisons. (**B**) Identification of common and specific DMRs across ES-C, ES-EA and EA-C comparison in CG and CHG context. DMRs common between ES-C and EA-C comparison and their methylation status are indicated in red. DMRs common between ES-C and ES-EA comparisons and their methylation are indicated in blue. Pie charts indicate the distribution of the commons DMRs associated with specific genomic features. C: Control leaf, ES: esca-plant symptomatic leaf, EA: esca-plant asymptomatic leaf

In the CG context, a similar number of DMRs were identified in ES-C (274) and ES-EA (275), with a majority being hypomethylated (ES-C: 84%, ES-EA: 86%). In contrast, only 55 CG-DMRs were found in the EA-C comparison, with a more balanced distribution (33 hypo-, 22 hyper-DMRs; Figure 3A). Ninety-three CG-DMRs were common between ES-C and ES-EA, of which 91 were hypomethylated. Only one of these was also present in the EA-C comparison (Figure 3B). CHH-DMRs were detected mainly in symptomatic tissues (11 in ES-C, 63 in ES-EA), with most being hypomethylated (ES-C: 91%, ES-EA: 92%; Figure 3A). Five CHH-DMRs were shared between ES-C and ES-EA, while none were found in EA-C (Figure S6). In the CHG context, a comparable number of DMRs were identified in ES-EA (225) and EA-C (247), but with opposite trends. ES-EA CHG-DMRs were predominantly hypomethylated (173 hypo-, 52 hyper-DMRs), while EA-C hyper-DMRs were prevalent (157 hyper-, 89 hypo-DMRs; Figure 3A). ES-C displayed the highest number of CHG-DMRs (384), with a near-balanced distribution (212 hypo-, 172 hyper-DMRs; Figure 3A). A large number of CHG-DMRs were shared between ES-C and EA-C (110; 69 hyper, 41 hypo-DMRs; Figure 3B), while fewer were common to ES-C and ES-EA (40; 39 hypo-, 1 hyper-DMRs; Figure 3B).

Considering all comparisons, approximately half of the DMRs were associated with transposable elements (TEs) (EA-C: 140/303; ES-C: 310/669; ES-EA: 299/563; Figure S7). TE-DMRs located in intergenic regions were more prevalent abundant in symptomatic tissues (ES-C: 47%, ES-EA: 56%) than in asymptomatic leaves (EA-C: 31%; Figure S6). Similarly, TE-DMRs in promoter regions were more frequent in symptomatic tissues (ES-C: 17%, ES-EA: 24%) than in EA-C (11%) (Figure S7).

We further examined DMRs located within gene bodies (intragenic-DMRs) and in promoters (2kb upstream of gene start; prom-DMRs). The total number of gene-associated DMRs (gene-DMRs = intragenic + prom-DMRs) was higher in comparisons with ES leaves (ES-C: 386, ES-EA: 303) than in EA-C (211) (Figure S7). Prom-DMRs accounted for a greater proportion of gene-DMRs in comparisons with ES leaves (ES-EA: 31%, ES-C: 42%) than with EA leaves (EA-C: 16%) (Figure S7).

Among the gene-DMRs common between ES-C and ES-EA, nearly half of the CG-DMRs (48/93) and CHG-DMRs (22/40) overlapped with gene regions, including a substantial number of TE-associated promoter DMRs (CG: 18/48, CHG: 11/22) (Figure 3B). In contrast, of the 23 CG-DMRs shared between ES-C and EA-C, only 6 were gene-DMRs. However, a large proportion of the 110 CHG-DMRs common between ES-C and EA-C were gene-associated (78/110), many of which were located within gene bodies, particularly introns (27 TE-intron DMRs, and 27 intron DMRs; Figure 3B).

These findings reveal an extensive methylome reprogramming in both symptomatic and asymptomatic tissues of esca-affected plants, with distinct methylation patterns across sequence contexts and genomic features.

### DNA methylation changes and gene activation can be associated in symptomatic leaves

We assessed the relations between gene expression and DNA methylation by analyzing the co-location between DMRs and DEGs in the ES-EA and ES-C analyses to identify DEGs associated with at least one DMR. These are referred to as differentially methylated and expressed genes (DMEGs). We used either the coordinate of the gene body (DMEGs; Table 1), or of the promoter region (2kb upstream gene start: prom-DMEGs; Table 2). In total, 35 DMEGs (15 in ES-C, 13 in ES-EA, and 7 common to both datasets), and 32 prom-DMEGs (13 in ES-C, 12 in ES-EA, and 7 common to both datasets) were identified. Of note, 73% (16 out of 22) and 85% (17 out of 20) DMEGs of the ES-C and ES-EA comparisons respectively, were hypo-methylated in ES leaves (Figure 4A, Table 1). Similarly, all prom-DMEGs were hypo-methylated in ES compared to EA and C leaves (Figure 4A, Table 2). In some cases, a single DEG contains multiple DMRs that in all cases show the same trend of methylation (Table 1 and 2). Among these, we identified genes containing hypo-methylated DMRs localized in both the promoter and gene body comparing ES-C (*Vitvi19g01872*, uncharacterized protein) and ES-EA (Vitvi16g04349, Stilbene/Resveratrol synthase 3) (Table 2).

**Figure 4.**
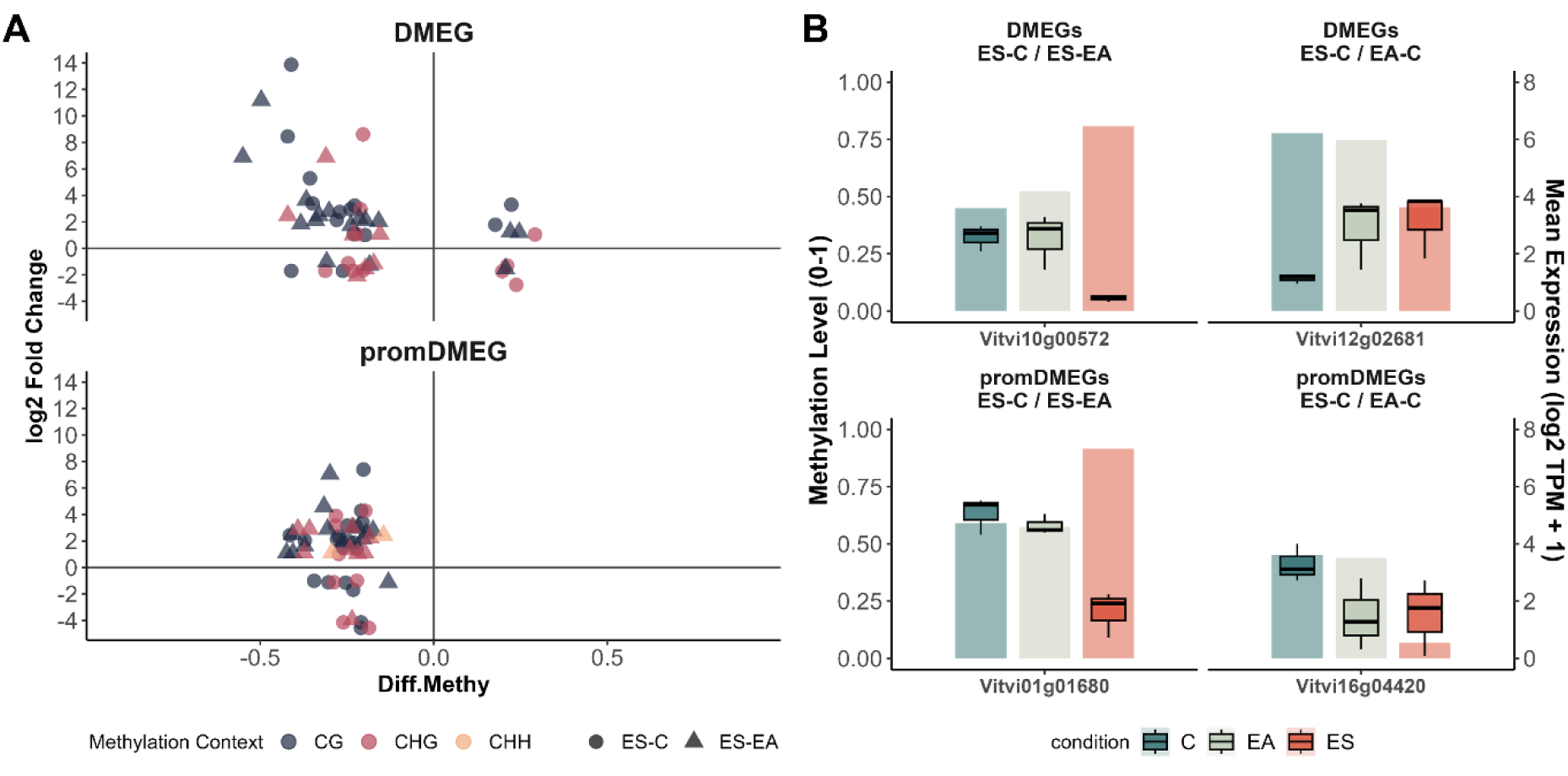
Some genes exhibit differential expression associated with changes in DNA-methylation. (**A**) Scatter plots represent the differential expression of genes in the ES-C (circles) and ES-EA (triangles) analyses. Each dot represents a differentially methylated region (DMR) associated with a differentially expressed gene (DEG). The position of each dot on the graph is defined by the differential expression value of a gene (Log2 Fold Change) on the y-axis, and the differential methylation value (Diff.Methy) of the associated DMR on the x-axis. DEGs associated with DMR localized within the gene (DMEG) are separated from those associated with DMRs localized at gene promoter (promDMEGs). (**B**) Boxplots represent the distribution of DNA methylation levels (0-1) across samples for two (prom)DMEGs common between ES-C and ES-EA analysis (ES-C/ES-EA) and common between ES-C and EA-C analysis (ES-C/EA-C). Bar heights indicate the corresponding mean gene expression levels (log2 TPM+1). C: Control leaf, ES: esca-plant symptomatic leaf, EA: esca-plant asymptomatic leaf

**Table 1.**
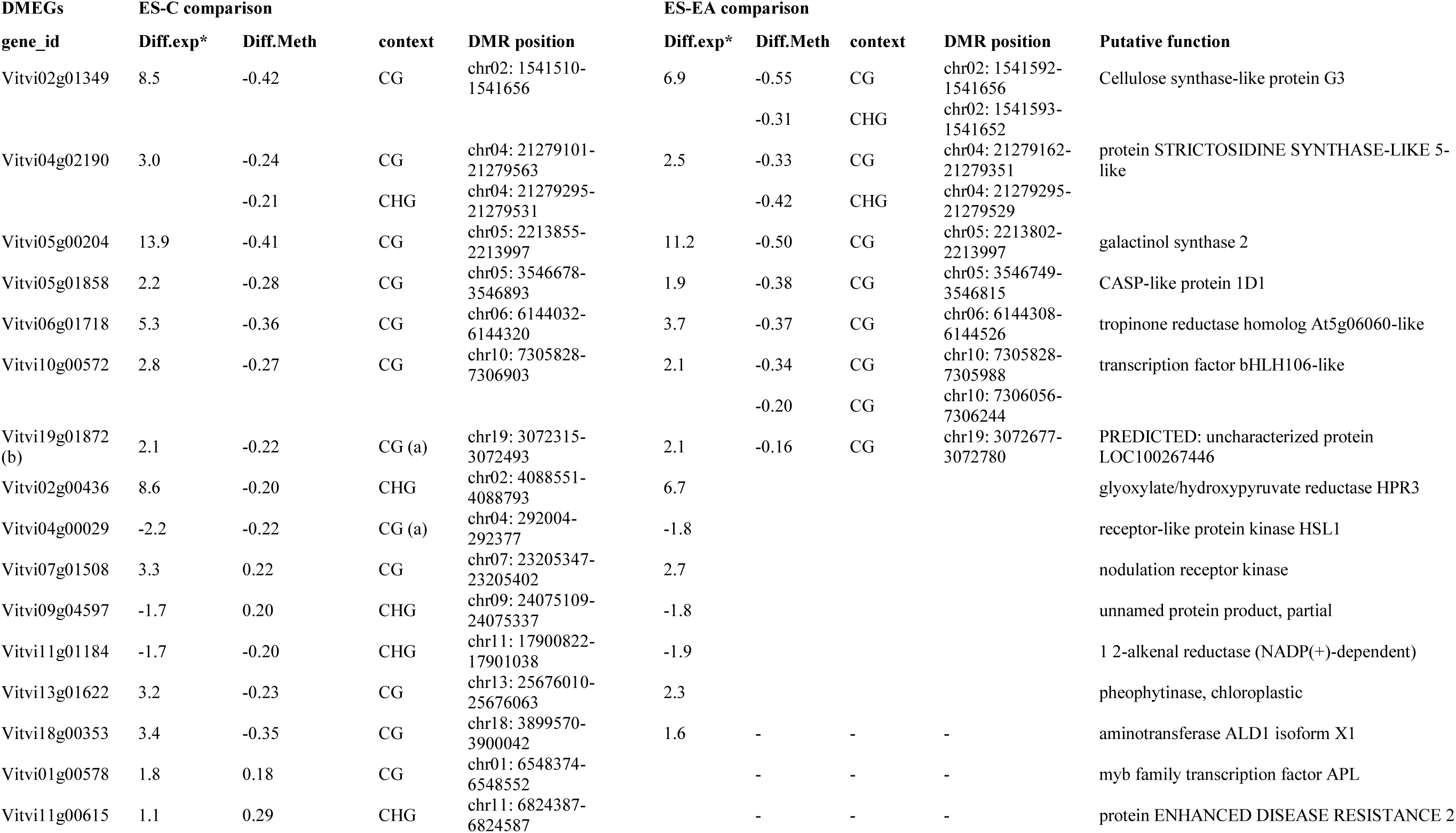
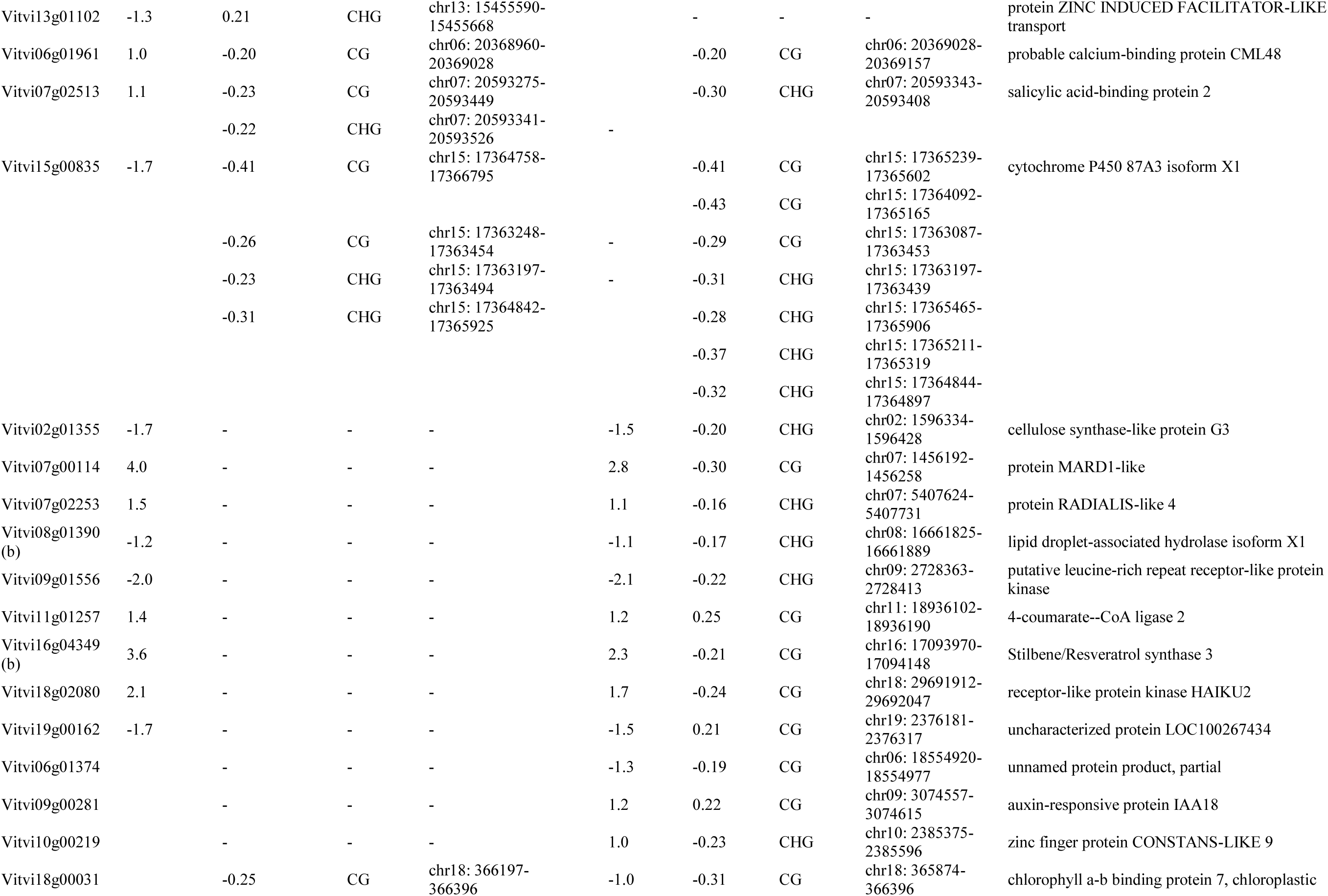
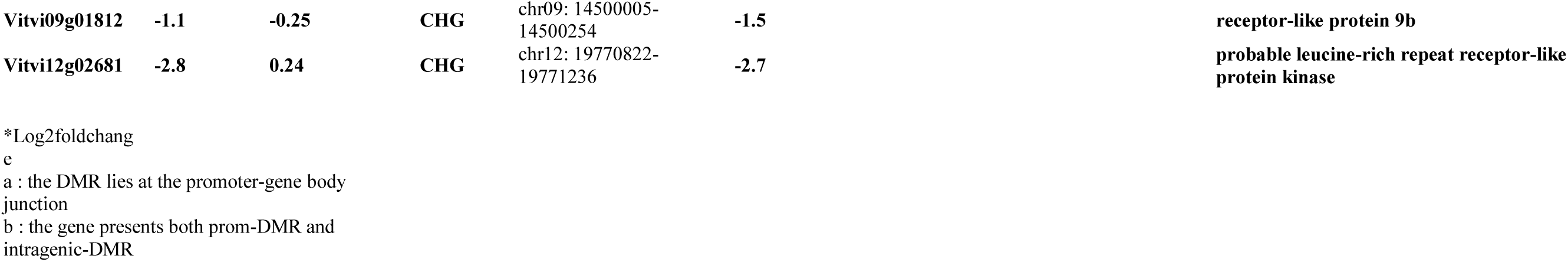
Differentially Methylated and Expressed Genes (DMEGs) and their associated DMRs identified in the ES-C and ES-EA comparisons. The table reports differential methylation (Diff.Meth) and gene expression 2fold change (Diff. exp) values observed for each genes in each comparison. Diff.Meth: positive values represent hypermethylation (+), negative values represent hypomethylation (-); Diff.exp: positive values represent up-regulation, negative values represent dow-regulation.

**Table 2.**
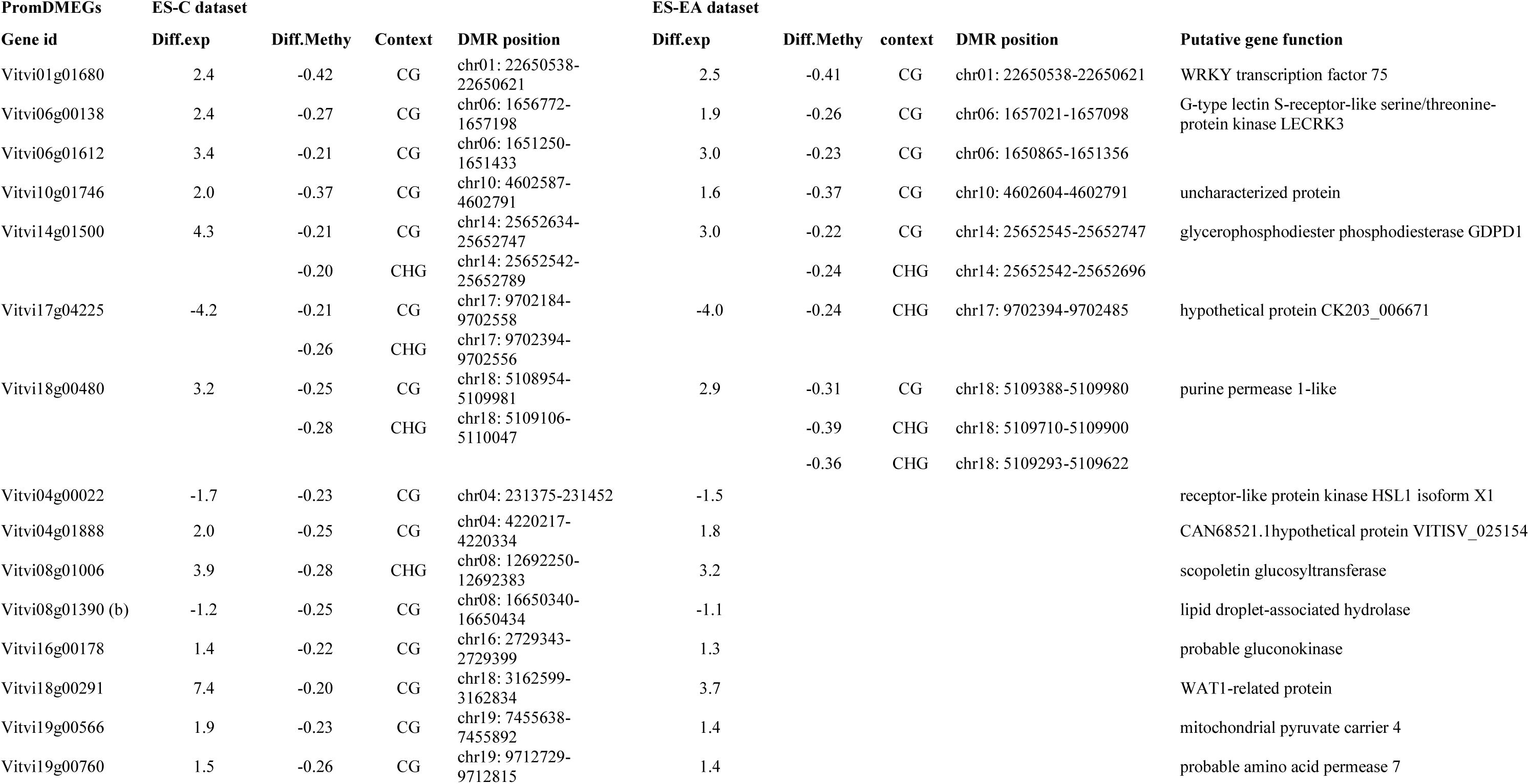
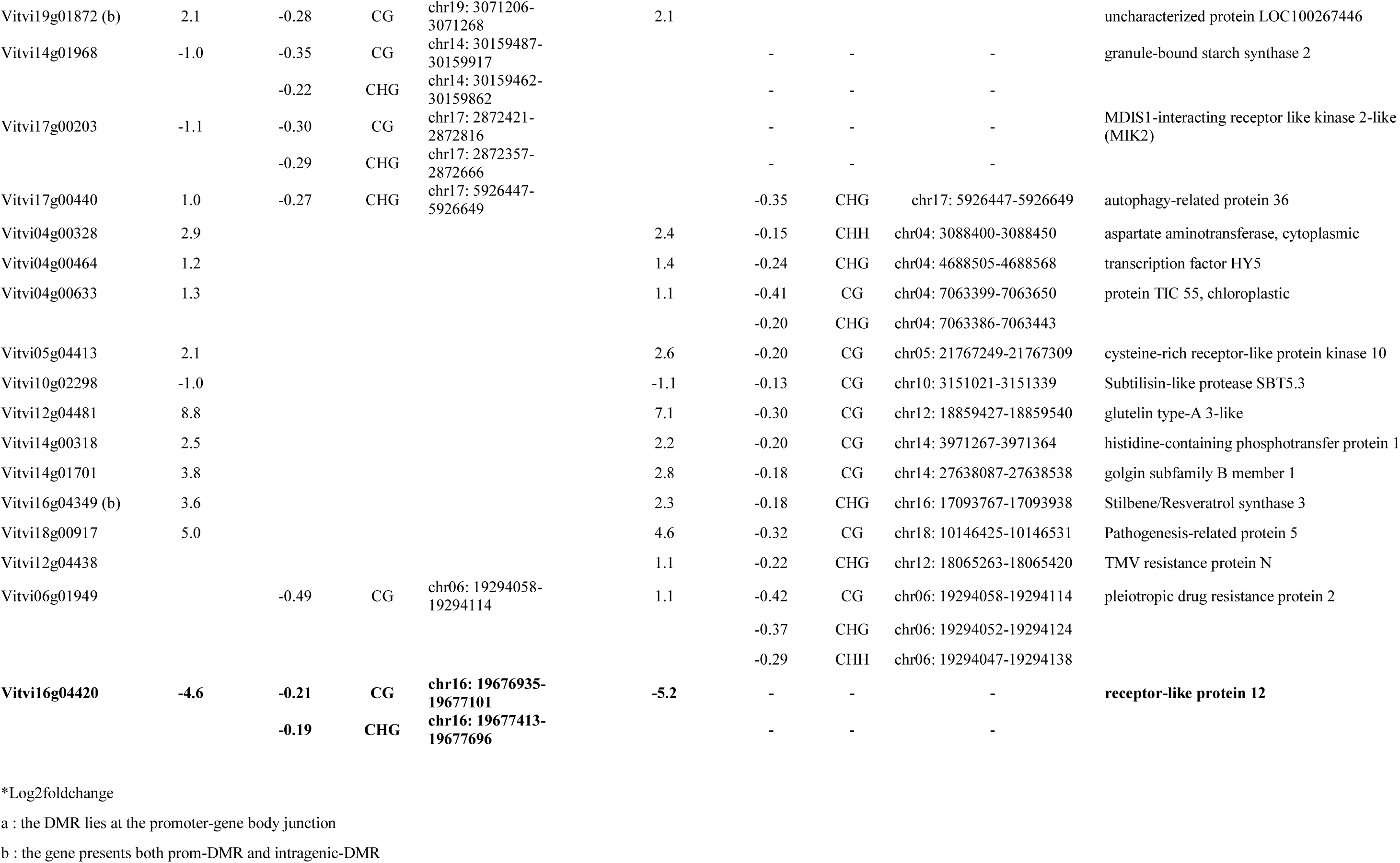
Differentially Methylated promoter of Differentially Expressed Genes (promDMEGs) and their associated DMRs identified in the ES-C and ES-EA comparisons. The table reports differential methylation (Diff.Meth) and gene expression 2fold change (Diff. exp) values observed at the 2kb region flankin Transcription Start Site for each genes in each comparison. Diff.Meth: positive values represent hypermethylation (+), negative values represent hypomethylation (-); Diff.exp: positive values represent up-regulation, negative values represent dow-regulation. C: Control leaf, ES: esca-plant symptomatic leaf, EA: esca-plant asymptomatic leaf

When considering the relationship between DNA methylation dynamic and gene expression, the most frequent pattern is the association of Up-DEGs with hypo-DMRs (18 of the 35 DMEGs and 25 of the 32 prom-DMEGs) identified in ES leaves compared to non-symptomatic leaves (EA and C, respectively) (Figure 4, Table 1, 2). These include the 7 DMEGs common to the ES-C and ES-EA comparisons, that encode a cellulose synthase (*Vitvi02g01349*), a Strictosidine synthase (*Vitvi04g02190*), a galactinol synthase 2 (*Vitvi05g00204*), a CASP-like protein 1D1 (*Vitvi05g01858*), a tropinone reductase (*Vitvi06g01718*), and a transcription factor, bHLH106 (*Vitvi10g00572*) (Figure 4B, Table 1). A similar pattern was found in 6 of the 7 shared prom-DMEGs, including those that encode a WRKY transcription factor (*Vitvi01g01680*), a G-type lectin-S-receptor like serine/threonin-protein kinases LECRK3 (*Vitvi06g00138*, *Vitvi06g01612*), a glycerophosphodiester phosphodiesterase GDPD1 (*Vitvi14g01500*) and a purine permease (*Vitvi18g00480*), and only one was hypomethylated and downregulated (*Vitvi17g04225,* hypothetical protein CK203) (Figure 4B, Table 2).

Among the 84 gene-DMRs common between the ES-C and EA-C datasets (Figure 3B), three were associated with differential gene expression in the ES-C comparisons: two DMEGs (*Vitvi09g01812* and *Vitvi12g02681*) and one promDMEG (*Vitvi16g04420*), all encoding receptor-like proteins (Table 3). ES and EA leaves displayed similar methylation pattern at these loci: *Vitvi09g01812* and *Vitvi16g04420* were hypomethylated, while *Vitvi12g02681* was hypermethylated. However, downregulation of these genes was only observed in ES leaves (Figure 4B, Table 3).

**Table 3.** Genes exhibiting common methylation patterns in ES and EA leaves as compared to C, but differential expression only in ES samples. C: Control leaf, ES: esca-plant symptomatic leaf, EA: esca-plant asymptomatic leaf

| Gene | Diff.Exp* | DMR type | DMR position | Diff.Meth | Context | Dataset | Gene function |
| --- | --- | --- | --- | --- | --- | --- | --- |
| Vitvi09g01812 | -1.1 | DMG | chr09:14500005-14500254 | -0.25 | CHG | ES-C | Receptor-like protein 9b |
|  | NA | DMG | chr09:14500005-14500254 | -0.26 | CHG | EA-C |  |
|  | -1.5 | NA | NA | NA | NA | ES-EA |  |
| Vitvi12g02681 | -2.8 | DMG | chr12:19770822-19771236 | 0.24 | CHG | ES-C | Probable leucine-rich repeat receptor-like protein |
|  | NA | DMG | chr12:19770822-19771092 | 0.24 | CHG | EA-C |  |
|  | -2.7 | NA | NA | NA | NA | ES-EA |  |
| Vitvi16g04420 | -4.6 | prom-DMR | chr16:19676935-19677101 | -0.21 | CG | ES-C | Receptor-like protein 12 |
|  |  | prom-DMR | chr16:19677413-19677696 | -0.19 | CHG |  |  |
|  | NA | prom-DMR | chr16:19676935-19677101 | -0.21 | CG | EA-C |  |
|  | NA | prom-DMR | chr16:19677568-19677633 | -0.16 | CHG |  |  |
|  | -5.2 | NA | NA | NA | NA | ES-EA |  |

\*Log2FoldChange

Together, these results show that (prom-)DMEGs are enriched in hypomethylated regions associated with increased genes expression in symptomatic leaves (ES) compared to asymptomatic leaves (EA and C). Some of these methylation changes were also detected in asymptomatic sector of esca-plants (EA), but without corresponding changes in gene expression.

### Esca-plants display specific methylation patterns before the symptom development

We performed methylome analysis using an independent set of grapevines affected by esca. We leveraged the experimental system described in Bortolami et al. (2021), that involves potted plants that were transplanted from the field to the greenhouse in 2018 (Figure S1). These plants had either exhibited esca foliar symptoms (pE) or remained asymptomatic (pC) between 2012 and 2017. Greenhouse plants were sampled early in the 2018 and 2019 growing seasons, before any foliar symptoms were observed (Figure S1). Two plant types were selected for WGBS methylome profiling (Table S3): symptomatic plants that had exhibited Esca symptom both in the field and for two consecutive years after transplantation, and asymptomatic plants that remained symptom-free throughout the study (Table S1).

We first analyzed methylation differences between pE and pC plants separately in 2018 and 2019. A comparable number of DMRs was identified in both years, with 348 DMRs in 2018 and 304 in 2019 (Figure S8A). However, the methylation patterns differed notably. In 2018, hyper-methylated DMRs accounted for 59.9% of the total DMRs identified (208 of 347), whereas in 2019, they represented only 37.2% (113 of 304) (Figure S8B). This change is primarily driven by a marked shift of DMR direction in the CHG context: in 2018, CHG DMRs were predominantly hyper-methylated (160 hyper- vs. 112 hypo-DMRs), while in 2019, this trend was reversed, with a majority of hypo-methylated CHG DMRs (162 hypo- vs. 70 hypermethylated) (Figure S8A).

When examining the genomic distribution of DMRs, we found that approximately 41% (2018) and 36% (2019) were associated with TEs. Hence, the majority of DMRs were located within genes (197 DMGs in 2018; 165 DMGs in 2019) or in promoter regions (41 promDMRs in 2018; 29 in 2019), together accounting for 69% and 64% of all DMRs identified in 2018 and 2019, respectively (Figure S8B, S9).

These results indicate that plants expressing esca symptoms during multiple years display distinct methylation patterns compared to controls, even before symptoms can be detected on leaves.

### Toward the identification of specific methylation patterns of Esca-plants

To pinpoint DNA methylation signatures specifically associated with plants developing symptoms of the esca infection, we aimed to identify DMRs that are maintained independently of the plant growing conditions and symptom development, but would differentiate any sectors of plants with symptoms (ES; EA, pE), from those not developing symptoms (C, pC).

The comparison by context of the DMRs distinguishing pE and pC plant in both 2018 and 2019 with those differentiating symptomatic from control plants in the vineyard experiment (ES-C and EA-C common DMRs; Figure 5A) identified 3 DMRs consistently associated with esca symptoms. To further explore whether these loci are part of broader methylation hotspots, all DMRs previously identified in the ES-C, EA-C, pE-pC 2018, and pE-pC 2019 comparison, regardless of methylation context, were mapped onto the grapevine genome (Methods, Figure S10A). This analysis revealed three genomic regions of interest, each characterized by the local accumulation of overlapping DMRs. Notably, these regions coincide with the three DMRs identified in the cross-comparison analysis, reinforcing their relevance as loci of disease-associated methylation change (Figure S10B, Table 4). Two of these regions (regions 66 and 68) are located on chromosome 13, and one (region 9) on chromosome 17. Region 68 include CG- and CHG-DMRs from all comparisons except pE-pC 2018, while regions 66 and 6 were only associated with CHG-DMRs (Figure S10B, Table 4).

**Figure 5.**
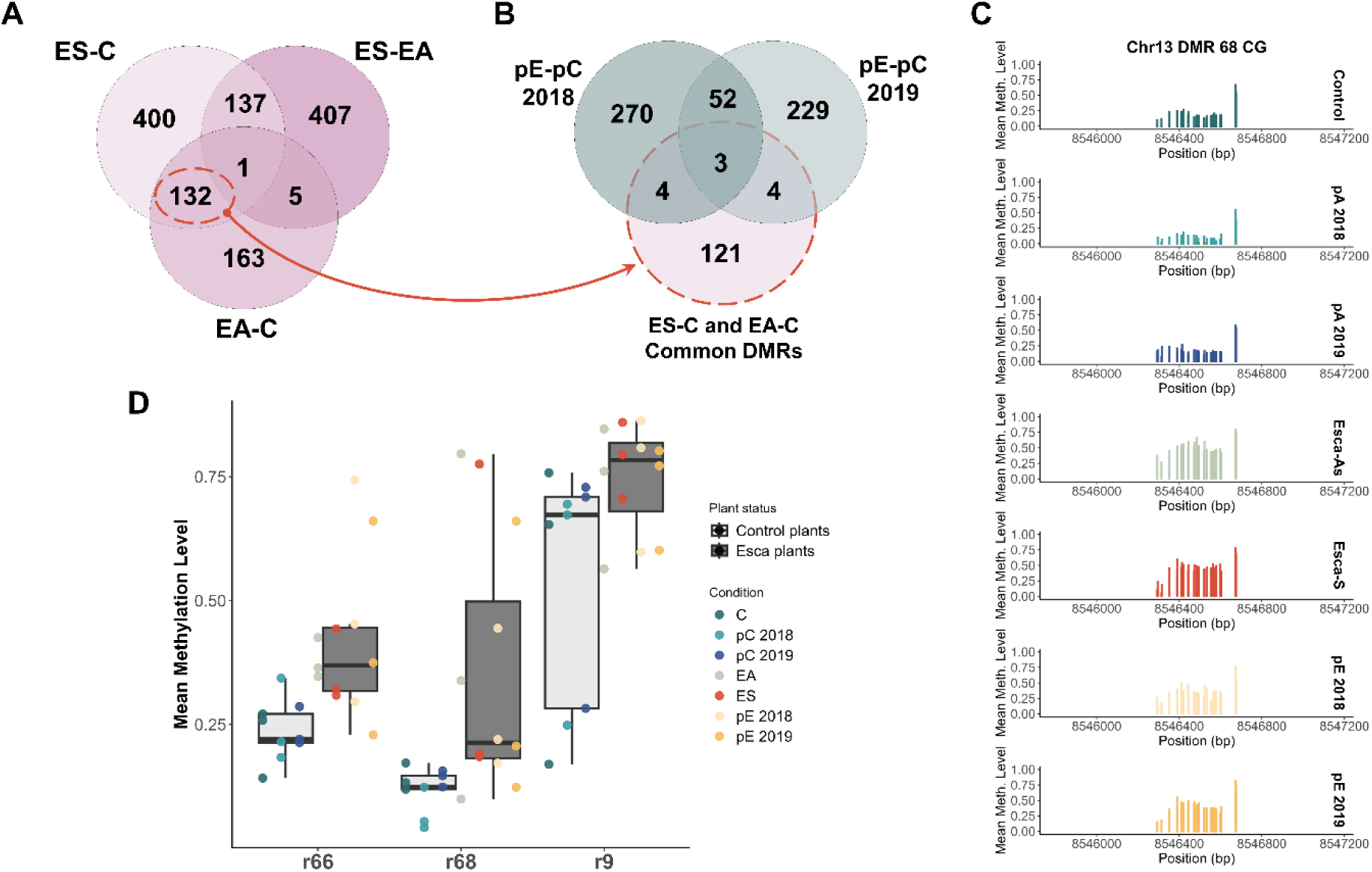
Identification of conserved differentially methylated regions in esca-plants across vineyard and greenhouse experiments. (A) Venn diagram showing the proportion of specific and common DMRs identified in the ES-C, ES-EA and EA-C datasets. (B) Venn diagram comparing the common DMRs in EA-C and ES-C to the DMRs that differentiated pE and pC plants in 2018 and 2019, before symptoms developed. (C) Details of CG methylation levels and distribution over the DMR region 68. Each bar represent the average methylation observed in three replicate samples at each cytosine position. (D) Boxplot representing the methylation levels for each regions in leaves sampled from control plants (light shade;C, pC) and esca plants (dark shade; ES, EA, pE). Samples are presented by colored dots as indicated. C: Control leaf, ES: esca-plant symptomatic leaf, EA: esca-plant asymptomatic leaf

**Table 4.**
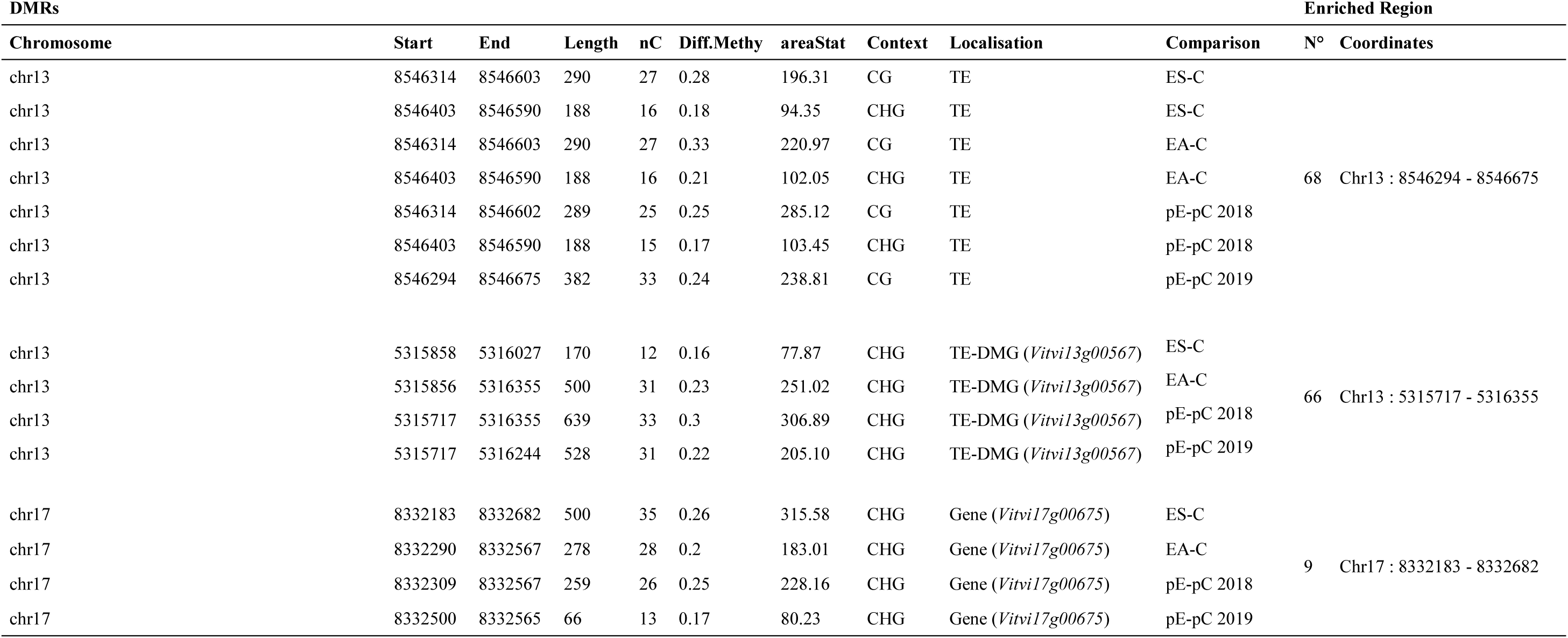
Details about the DMRs shared between the ES-C, EA-C, and pE-pC datasets, as well as the three enriched regions identified, are provided.

We further examined the methylation levels in regions 68, 66 and 9. All DMRs had higher methylation levels in esca-plants (23%, 20% and 17% higher mean methylation in regions 68, 9 and 66 respectively) than in control plants (Figure 5C,D, Figure S11-13, Table S4).

However, variation in methylation levels were observed within conditions, depending on the status of the origin of the plant (Figure 5D, Figure S11-13, Table S4-5). In region 68, ES, EA and pE samples showed important methylation levels variations, ranging from 10% to 78%, while control DNA methylation levels varied much less, ranging from 4% to 17% (Figure S11, Table S5). In contrast, region 9 displayed an opposite trend: esca-plants consistently have a methylation levels above 60% (60-86%) whereas the methylation level of control samples ranged from 17% to 76% (Figure S12, Table S5).

Overall, these findings reveal that the recurring expression of esca-symptoms in plants is associated with changes in DNA methylation at specific loci, independently of growing conditions and of foliar symptom expression.

## Discussion

Esca is a major grapevine trunk disease that significantly reduces its productivity and longevity. Despite the dramatic consequences of the esca disease in viticulture, the underlying molecular mechanisms remain poorly understood. In the present study we aimed at generating a comprehensive view of the plant response to esca. To this end, we took advantage of the very specific situation of plants affected by esca, which can present sectors developing symptoms while some other do not. We have therefore compared the transcriptomes and methylomes of symptomatic (ES) and asymptomatic (EA) leaves from esca-plants and of leaves of control-plants that have not expressed symptoms during the 5 years that preceded the experiment (C). In addition, as symptoms developed late during the growing season, we also analyzed the methylation profiles of plants before any foliar symptoms can be detected. Overall, our results show that metabolic changes and transcriptomic reprogramming were restricted to symptomatic leaves and partially associated with local changes in DNA methylation. However, asymptomatic leaves presented distinct DNA methylation, some of which were also observed in symptomatic tissues, suggesting a systemic response to esca at the epigenetic level. Notably, a subset of these epigenetic marks were observable prior to symptom emergence, highlighting their potential as early biomarkers for esca-disease detection in grapevine.

### Metabolic and transcriptomic response ton esca are restricted to symptomatic sectors

Recent studies have highlighted significant metabolic reprogramming in esca-symptomatic grapevines, including alterations in carbohydrate metabolism and the accumulation of secondary metabolites (Magnin-Robert *et al*., 2016; Bortolami *et al*., 2021; Weiller *et al*., 2024). However, these analyses have focused exclusively on symptomatic tissues from symptomatic vines without assessing asymptomatic sectors developing on symptomatic plants. In our study, by analyzing asymptomatic leaves collected from plants expressing esca foliar symptoms, we provide evidence that changes in primary metabolite assumulation, particularly carbohydrates, might be restricted to symptomatic canopy sectors (Figure 1B, C).

There is growing evidence that gene expression changes associated with Esca symptom expression may, to some extent, reflect these metabolic changes. This is supported by previous studies reporting altered expression of genes involved in stress responses and primary and secondary metabolism in the wood and annual shoots of symptomatic grapevines (Magnin-Robert *et al*., 2016). More recently, transcriptomic analyses of wood tissues from vines inoculated with esca-associated pathogens revealed extensive transcriptional reprogramming, particularly affecting stress-related and metabolic pathways (Romeo-Oliván *et al*., 2024). However, transcriptomic investigations have so far focused exclusively on woody tissues, and no comparable analyses have been conducted on leaves of symptomatic and asymptomatic sectors of a same affected plant.

To address this gap, we investigated transcriptomic changes in leaves of esca-plants using RNA-seq. Our results demonstrate that symptomatic leaves (ES) differ in similar ways from non-symptomatic leaves, whether sampled from the same symptomatic plant (EA), or from asymptomatic control-plants (C). Notably, a similar trend was previously observed in green stems, where genes related to stress and pathogen responses were similarly expressed in asymptomatic tissues of esca and control plants (Magnin-Robert *et al*., 2016). In agreement with this, a comparable number of DEGs were identified in the ES-EA and ES-C comparisons, with a large majority being shared between both. This high degree of overlap suggests that transcriptional reprogramming is primarily restricted to symptomatic tissues, while asymptomatic leaves regardless of their origin remain largely similar at the transcriptomic level (Figure 2C). The DEGs affected include genes related to transport, transcriptional regulation, and biotic stress responses that are up-regulated while those related to photosynthesis and growth are down-regulated (Figure 2D). Hence, both metabolic modifications and transcriptome remodeling are associated with the development of symptoms in esca-plants, while non-symptomatic leaves of esca-plants do not differ significantly from leaves of control plants.

### The expression of Esca-symptoms triggers local and systemic epigenetic changes in symptomatic and asymptomatic canopy sectors

Epigenetic information has recently been shown as a major regulatory process involved in the response of plants to both abiotic and biotic stresses (Zhang, Lang and Zhu, 2018) (Lämke and Bäurle, 2017; Zhang, Lang and Zhu, 2018). An important aspect of the response of plants to pathogen attacks concern the remodeling of the DNA methylation patterns that are deeply influenced by the type of pathogen and the time of analysis (Arora *et al*., 2022). Current results show that hypomethylation mediates a better tolerance to biotrophic pathogens while hypermethylation has an opposite effect. Interestingly necrotrophic pathogens behave in an opposite way. Altogether, these data highlight the relevance of analyzing DNA methylation of specific plant-pathogen interactions (Hewezi *et al*., 2018; Alonso, Ramos-Cruz and Becker, 2019). However, esca is peculiar as it does not rely on one pathogen but on a highly complex consortium of different microorganisms that is difficult to fully define (Larignon and Dubos, 1997; Del Frari *et al*., 2019). Furthermore, their presence is not necessarily associated with the development of symptoms in a specific tissue, and evidence suggest that symptoms may result from a combination of fungi toxins and physiological effects related to abiotic stresses (Bortolami *et al*., 2019; Lecomte *et al*., 2024). This makes the study of esca disease challenging and so far there is no data available on a potential epigenetic reprogramming of plants developing esca symptoms, unlike many other patho-systems in plants (Dowen *et al*., 2012; Hewezi *et al*., 2018; DiBiase *et al*., 2024).

Our results identified several DMRs associated with changes in DNA methylation in ES leaves were identified (Figure 3), consistent with a specific epigenetic remodeling associated with the development of leaf-symptoms. The large majority of these DMRs were hypomethylated in ES leaves in the CG and CHH contexts. In addition, some of these DMRs were common between ES-C and ES-EA comparison (Figure 3B, Figure S6) supporting the idea that these methylation changes represent local consequences of symptom development. Regarding CHG-DMRs, their numbers and patterns differed depending on the asymptomatic leaf considered in the comparison. While CHG-DMRs identified in ES-EA displayed a clear biased toward hypomethylation, those identified in ES-C were equivalently distributed between hypo- and hyper-DMRs, highlighting difference in methylation patterns between asymptomatic leaves (EA and C) (Figure 3A).

The epigenetic landscapes of EA leaves (no symptoms, but sampled from symptomatic plants) diverged from those of leaves from control plants, although the number of DMRs identified was reduced by approximately two-fold as compared to DMRs found in ES-C and ES-EA comparisons. We observed that 133 DMRs differentiate both EA and ES leaves from controls (Figure 3B) suggesting a general effect of esca on grapevine leaf methylation levels, whether symptomatic or not. However, DMRs identified in the EA-C comparison revealed distinct methylation patterns. In particular, CHG-DMRs showed a notable bias toward hypermethylation (Figure 3A), a pattern contrasting with those observed when considering symptomatic leaves (ES-C and ES-EA). These observations suggests that in addition to a general effect of esca symptom expression on grapevine DNA methylation, the nature of these changes differs between symptomatic and asymptomatic sectors of a same symptomatic plant, potentially reflecting tissue-specific regulatory mechanisms.

To date, most studies investigating biotic stress-induced methylation changes in plants have focused on comparisons between inoculated and non-inoculated tissues, without distinguishing between inoculated and non-inoculated sectors within the same diseased individual (Dowen *et al*., 2012; Rambani *et al*., 2015; DiBiase *et al*., 2024). Taken together, our findings highlight the complex and spatially differentiated nature of DNA methylation changes in grapevine leaves during esca symptom expression. By considering symptomatic and asymptomatic tissues within the same plant, we were able to distinguish local methylation changes directly associated with foliar symptoms from systemic, symptom-independent effects. In particular, the divergent CHG methylation patterns across comparisons underline the context-specific nature of the response, suggesting that local and systemic cues may trigger distinct epigenetic changes. These results emphasize the importance of within-plant spatial resolution in methylome studies and open new perspectives for understanding how plants integrate local pathogen-induced signals with broader physiological responses.

### Transcriptomic changes observed in esca-symptomatic leaves are associated with epigenetic remodeling

The DMRs identified in ES samples showed a strong prevalence of hypo-methylation irrespective of sequence contexts, especially in CG (Figure 3A). In plants, DNA methylation can occur in promoter regions and gene bodies, with promoter methylation usually related to transcription inhibition, while demethylation may activate transcription (Zhang, Lang and Zhu, 2018). The functional importance of active promoter demethylation in regulating defense genes has been shown in *Arabidopsis* (Yu *et al*., 2013; Halter *et al*., 2021).

In ES leaves, we identified 25 hypomethylated and upregulated promDMEGs (Table 2). These included genes involved in signal transduction, such as two G-type lectin S receptor-like serine/threonine-protein kinases, LECRK3 (*Vitvi06g00138*, *Vitvi06g01612*), a gene family known in rice for enhancing biotic stress resistance (Liu *et al*., 2015), and a cysteine-rich receptor-like protein kinase (*Vitvi05g04413*), part of a family recognized as central regulators of plant signaling pathways, including biotic stress responses (Zeiner *et al*., 2023). We also identified genes encoding known pathogen-response protein such as TMV resistance protein N (*Vitvi12g04438*; (Whitham *et al*., 1994)) and Pathogenesis-Related Protein 5 (*Vitvi18g00917*; (Enoki and Suzuki, 2016), and other coding for transcription factors like HY5 (*Vitvi04g00464*), which regulates a broad range of downstream genes (Gangappa and Botto, 2016), and WRKY75, involved in plant stress responses (Shankar, Pandey and Pandey, 2013). Notably, recent work in tomato has shown that the activation of WRKY75, a transcription factor induced by biotic stress and a potential regulator of the jasmonic acid pathway, is associated with the presence of the activating histone mark H3K4me3 (histone H3 lysine 4 tri-methylation) at its promoter region (López-Galiano *et al*., 2018). This histone modification is known to promote active transcription and has also been characterized by facilitating active DNA de-methylation at transcription start sites of several genes in *Arabidopsis* (Wang *et al*., 2025).

Although recent studies have shown that intragenic cytosine methylation could regulate stress-responsive gene expression, such as in response to cold in Solanaceae (Xin *et al*., 2025), the precise functional role of gene body methylation remains under investigation (Bewick and Schmitz, 2017; Zhang, Lang and Zhu, 2018). In ES leaves, we identified 35 DMEGs, nearly half of which were hypomethylated and upregulated (Figure 4, Table 1). These 18 genes were associated with key biological processes involved in biotic stress responses, including cell wall remodeling (Cellulose synthase-like G3, *Vitvi02g01349*; (Wan *et al*., 2021; Hou *et al*., 2024), calcium signaling (CML48, *Vitvi06g0196*), pathogen response (Strictosidine synthase-like-5, *Vitvi04g02190*; (Sohani *et al*., 2009), secondary metabolism (Stilbene/Resveratrol Synthase 3, *Vitvi16g04349*; (Vannozzi *et al*., 2012), disease resistance (ALD1, *Vitvi18g0035*; (Jiang *et al*., 2021), and transcriptional regulation (bHLH106, *Vitvi10g00572*; (Ahmad *et al*., 2015) (Table 1).

The consistent upregulation of these genes in ES samples suggest a potential role in the plant response to esca symptoms expression, particularly through the activation of defense mechanisms and stress-related pathways. Altogether, our findings suggest that DNA hypomethylation at specific loci may play a regulatory role in esca-induced transcriptional reprogramming in symptomatic tissues, similar to what has been observed in other plants such as *Arabidopsis* upon infection by *Pseudomonas syringae* (Dowen *et al*., 2012), or potato tuber exposed to *Fursarium* toxin deoxynivalenol (Shi *et al*., 2022). Furthermore, we identified 14 (prom)DMEGs that were common to both ES-EA and ES-C comparisons, suggesting a conserved and specific epigenetic signature of symptomatic leaves relative to asymptomatic tissues.

### Asymptomatic tissues reveal systemic methylation changes associated with esca

By comparing the methylation profiles of asymptomatic leaves, we identified 302 DMRs that differentiate EA from C leaves (Figure 3) suggesting that asymptomatic sectors of plants developing symptoms also undergo methylation changes. Among them, 169 DMRs are specific to EA leaves, and may reflect systemic effects associated to the formation of esca symptoms. The 132 DMRs were also found in the ES-C comparison, and are therefore not strictly associated with formation of symptoms. As such, they may rather reflect difference between esca-symptomatic plants and non-symptomatic ones (Figure 4-5).

Eighty-four of the 132 DMRs (C) (Supplementary data), were associated with genes, including genes linked to stress responses, such as *VASCULAR ASSOCIATED DEATH 1* (*Vitvi15g04322*) (Lorrain *et al*., 2004), a putative *RGA1* resistance gene (*Vitvi19g02270*) (Pathak *et al*., 2021), and *PHOS32* (*Vitvi10g00352*), a universal stress protein. We also detected transcription-related genes such as *POLYMERASE III SUBUNIT RPC5* (*Vitvi04g01256*) and an *RNA helicase DExH8* (*Vitvi07g00315*), as well as transcription factors including *TF-IIE* (*Vitvi11g01319*) and *TF-IIIB* (*Vitvi08g00112*). In addition, DMRs were found in genes involved in epigenetic regulation, including a *Histone-3-Leucine-79-methyltransferase* (*Vitvi14g01290*) and *HISTONE H2A.1* (*Vitvi10g04123*) (Zhou *et al*., 2015), and in signaling components like leucine-rich repeat (LRR) receptors (e.g, *Vitvi01g01434, Vitvi19g00052, Vitvi12g02681*) and receptor-like proteins (e.g., *Vitvi16g04420*, *Vitvi09g01812*) (Tang, Wang and Zhou, 2017). Interestingly, none of these genes were differentially expressed in EA leaves, but three of them were differentially expressed in ES compared to control leaves (Table 4). These 3 genes encore receptor proteins (Receptor-like protein 12*, Vitvi16g04420*; Receptor-like protein 9b, Vitvi09g01812 ; LRR-like protein kinase Vitvi12g02681).

The identification of common DMRs in asymptomatic and symptomatic leaves suggests that the expression of esca symptoms may trigger DNA methylation changes that prepare asymptomatic tissues to a rapid transcriptional activation upon pathogen detection, as previously proposed (Ramos-Cruz, Troyee and Becker, 2021). It is consistent with a systemic effect of Esca disease on the plant methylome, consistent with reports showing that pathogen-induced methylation alterations can occur in genomic regions distant from the infection site (Rodríguez-Negrete *et al*., 2013; Rambani *et al*., 2015).

### Toward the identification of stable methylation signatures of Esca-plants

By combining methylome data from vineyard-grown plants and those from plants transferred to the greenhouse, we have identified 3 DMRs that consistently distinguish plants that have expressed or are expressing Esca symptoms from those that have not. These DMRs may represent robust epigenetic markers of esca-plants. However, it is important to note that we observed inter-individual variability in methylation levels between samples. Three genomic regions corresponding to such conserved DMRs were identified. Two of them are located within gene bodies. One of them corresponds to a gene coding for a ZRANB3-like helicase (*Vitvi17g00675*), a protein known in mammals for its role in the maintenance of genome integrity (Ciccia *et al*., 2012). The other one codes for Beta-Taxilin (*Vitvi13g00567*), a pericentrosomal protein also described in human system that plays a role in protein quality control (McLendon *et al*., 2025) The function of these proteins is not described in plants so far, but they seem to be important to maintain the cellular homeostasis in mammals. Their potential role in esca disease remains to be elucidated.

However, the identification of these 3 genomic regions is consistent with an epigenetic signature of esca expression, potentially reflecting a systemic response and/or long-term imprints associated with disease development in the plant. Such methylation marks are possible candidates marking plants more likely to develop esca symptoms and could be used at early stage of the infection, much before any visible symptoms. As such, they may be used as early markers differentiating plants likely to develop symptoms of an esca infection from those that will not, and could offer a novel approach to screen for tolerant grapevines at the nursery stage, prior to planting. Validation of these candidate epigenetic markers now require further study involving a larger set of plants, and the comparison of their methylation profile before and during symptom expression. This approach would be critical to assess the dynamics of methylation at these loci, and to strengthen their relevance as predictive indicators of Esca susceptibility or progression in grapevine.

## Acknowledgements

We are grateful to the genotoul bioinformatics platform Toulouse Occitanie (Bioinfo Genotoul, https://doi.org/10.15454/1.5572369328961167E12) for providing advice, and giving access to computing facilities and storage resources.

This work was supported by the program ‘Plan National Dépérissement du Vignoble’ (FranceAgriMer/CNIV) in the framework of the projects EPIDEP (NUMERO) awarded to P.G. and PHYSIOPATH (22001150) awarded to C.E.L.D. M.B was in receipt of a grant financed by the program “Grand Projet de Recherche Bordeaux Plant Science” (TEPIMEMORY). B.R was in receipt of a grant financed by the program “PRIMA” (PROSIT).

## Author contributions

P.G and C.E.L.D conceived and designed the study. G.B, C.E.L.D and G.G were in charge of the greenhouse experiment. C.E.L.D and M.B contributed to vineyard sampling. M.B contributed to greenhouse sampling. M.B performed sample extraction, metabolite profiling, methylome and transcriptome analysis. V.G and B.R supervised NGS data analysis. M.B and P.G wrote the manuscript with input from all authors. P.G supervised the project.

## Data availability

The data for this study have been deposited in the European Nucleotide Archive (ENA) at EMBL-EBI under accession number PRJEB94866.

