## Supplementary Figures and Tables for "Esca Disease triggers local transcriptomic response and systemic DNA methylation changes in grapevine"

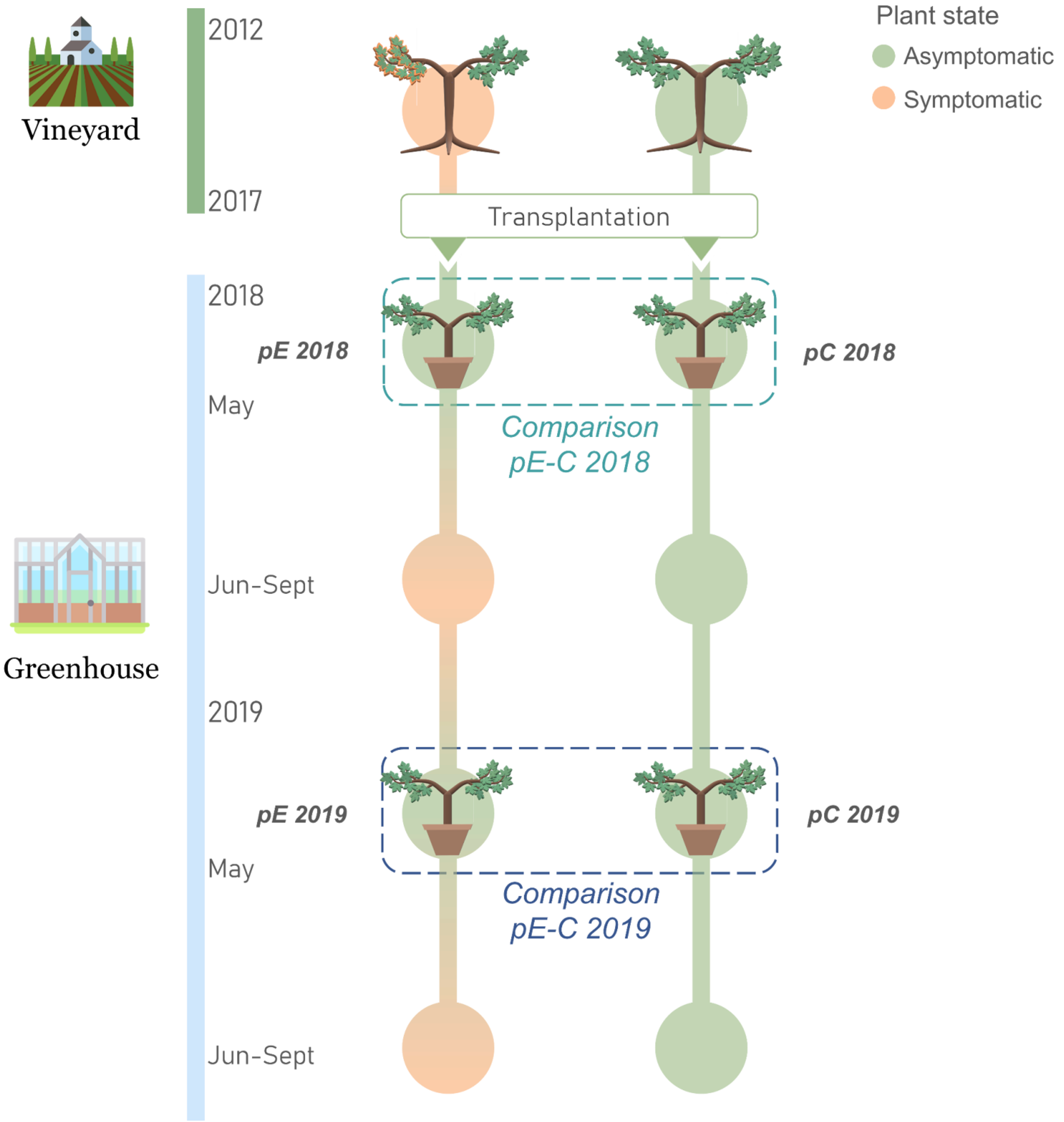

Figure S1. Design of the greenhouse experiment and sampling strategy: Plants were transplanted from the vineyard to the greenhouse in winter 2017. Leaves were sampled from the same plants at the beginning of 2018 and 2019 growing seasons before any leaf symptoms appeared. Samples were collected from plants expressing esca symptoms each year before and after transplantation (pE), and from plants that were asymptomatic before, and after transplantation (pC).

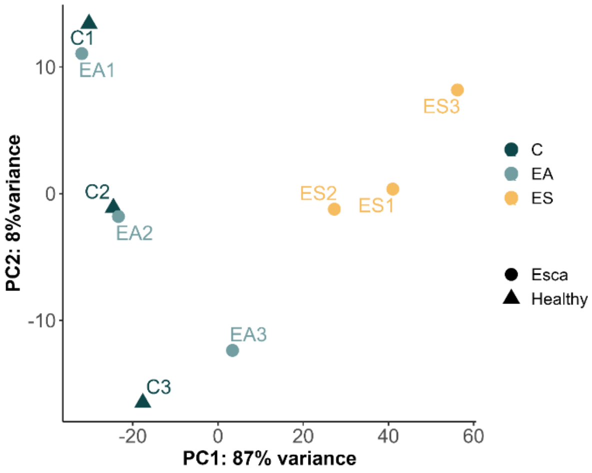

Figure S2. PCA of transcriptomic profiles of ES, EA and C leaves, based on all expressed genes.

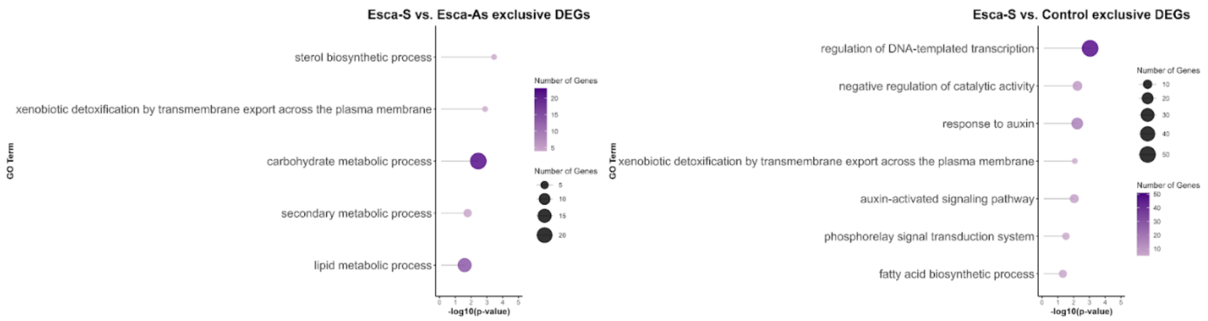

Figure S3. Gene Ontology analysis on ES-EA and ES-C specific DEGs.

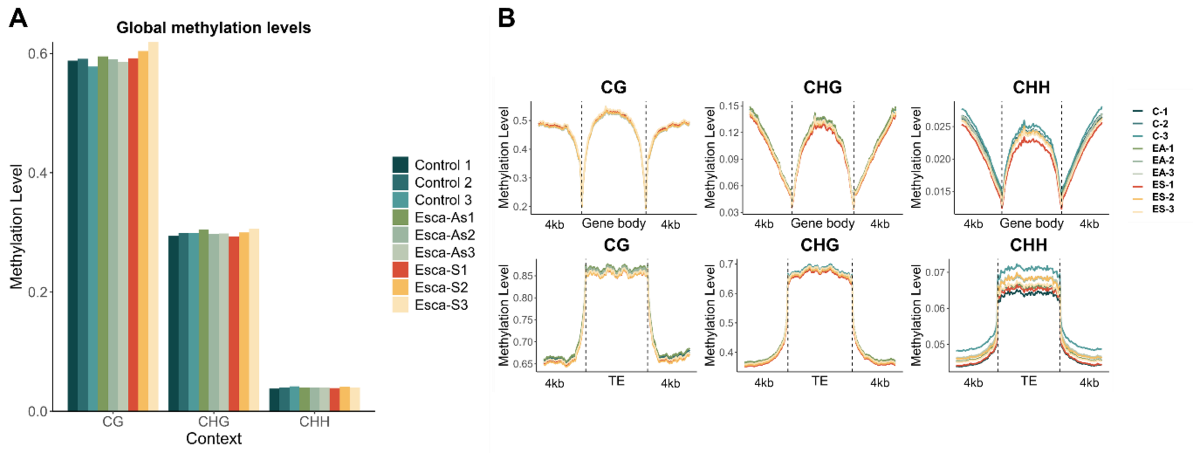

Figure S4. (**A**) Global methylation levels per sample in the three sequence contexts. (**B**) Distribution of DNA methylation along genes and transposable elements (TEs) in ES, EA and C leaves. (**C**)

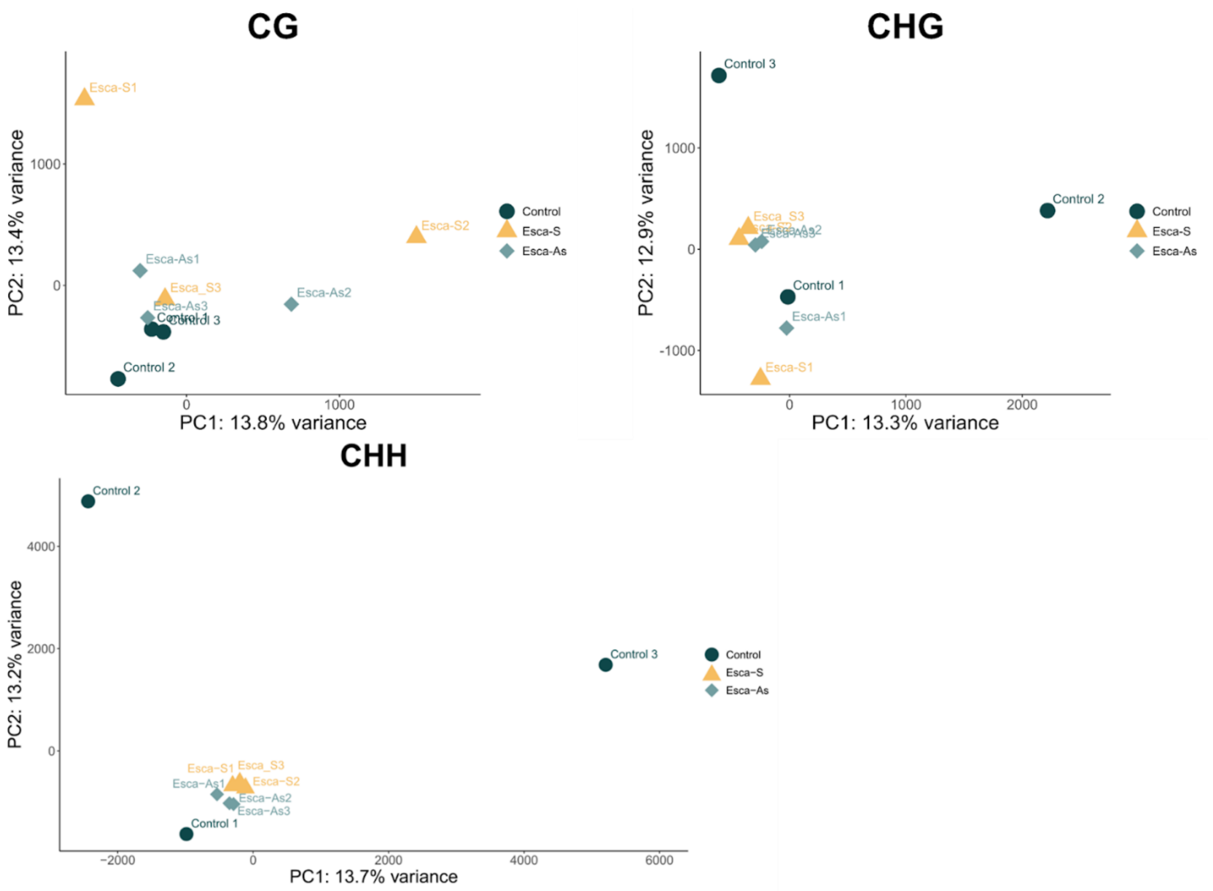

Figure S5. Principal Component analyses based on the global methylation profiles observed in each leaf samples collected in vineyard, in the three sequence contexts. Control: Control-plants, Esca-S: Esca-plants symptomatic leaf, Esca-AS: Esca-plants Asymptomatic leaf.

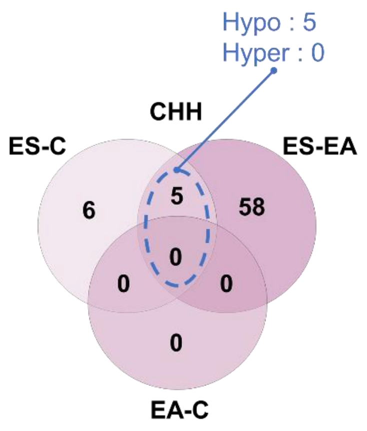

Figure S6. Identification of common and specific DMRs across ES-C, ES-EA and EA-C datasets in CHH context. DMRs shared between ES-C and ES-EA comparison and their methylation status are indicated in blue.

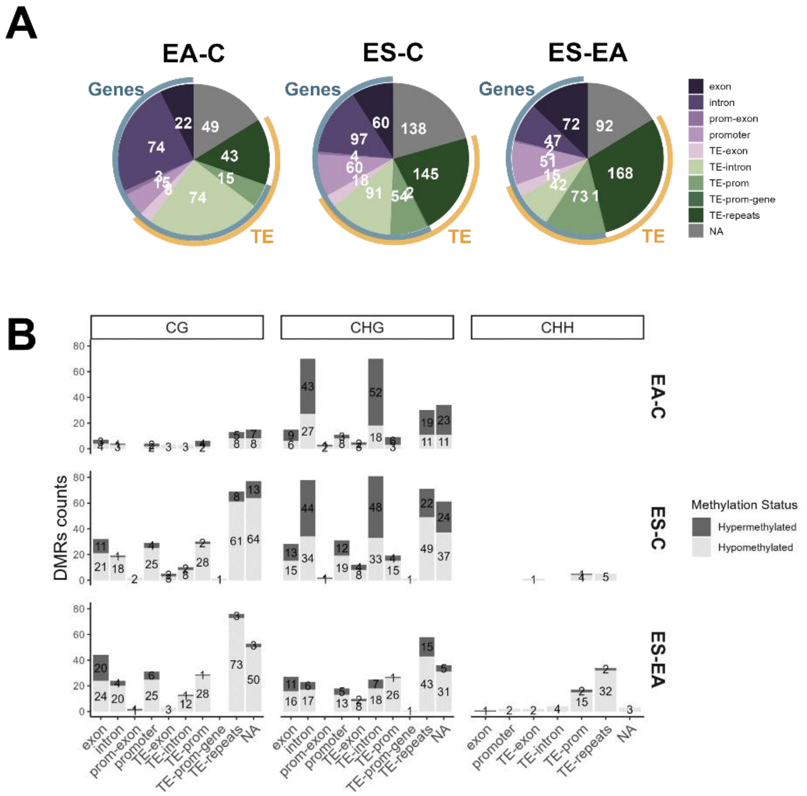

Figure S7. (**A**) Distribution of DMRs associated with specific genomic genomic features. The DMRs associated with TEs located in genes and promoter, as well as intergenic regions, are counted separately. The proportion of DMRs localized within genes or promoters is indicated by the inner-bar “Genes” and the proportion of DMRs localized within TE is indicated by the outer bas “TE”. (**B**) Number and methylation status of differentially methylated regions (DMRs) identified in EA-C, ES-C and ES-EA comparison according to their localization over specific genomic features.

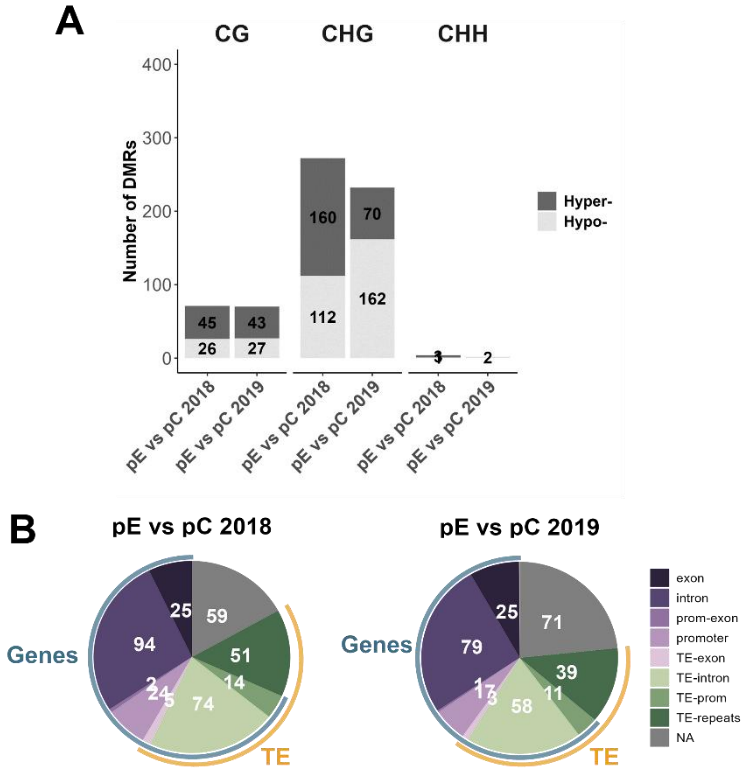

Figure S8. (**A**) Number and methylation status of differentially methylated regions (DMRs) identified in pE-C comparison in 2018 and 2019 (**B**) Distribution of DMRs associated with specific genomic features. The DMRs associated with TEs located in genes and promoters, as well as in intergenic regions are counted separately. The proportion of DMRs localized within genes or promoters is indicated by the inner bar "Genes", and the proportion of DMRs localized within TEs is indicated by the outer bar "TE.

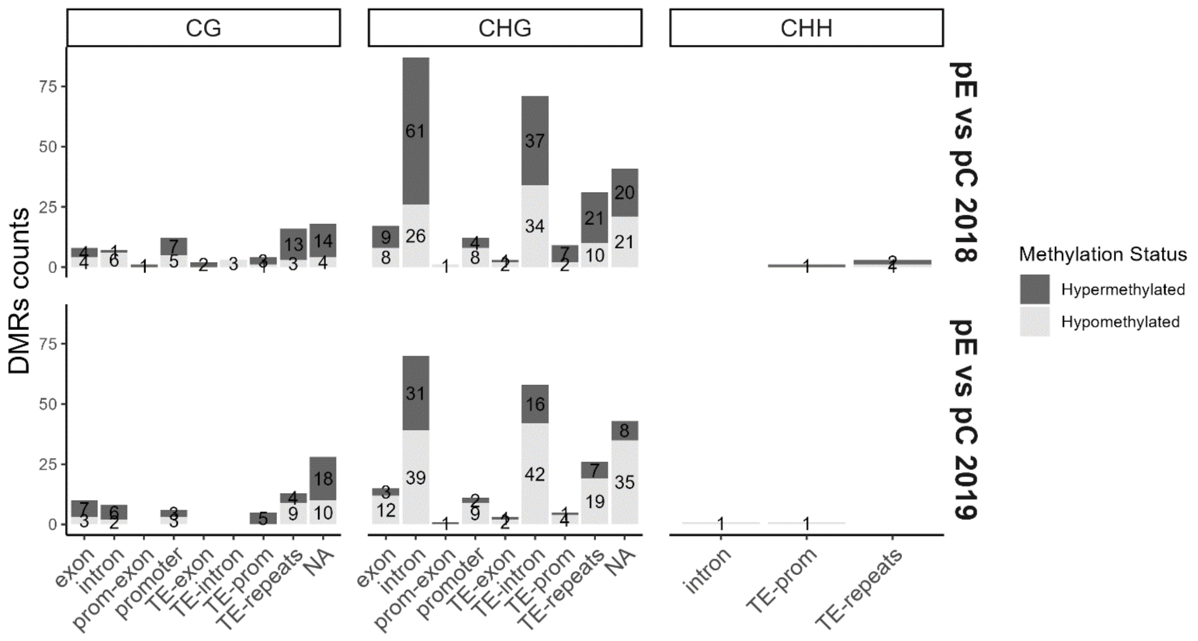

Figure S9. Number and methylation status of differentially methylated regions (DMRs) identified in pE-C 2018 and pE-C 2019 comparisons according to their localization over specific genomic features.

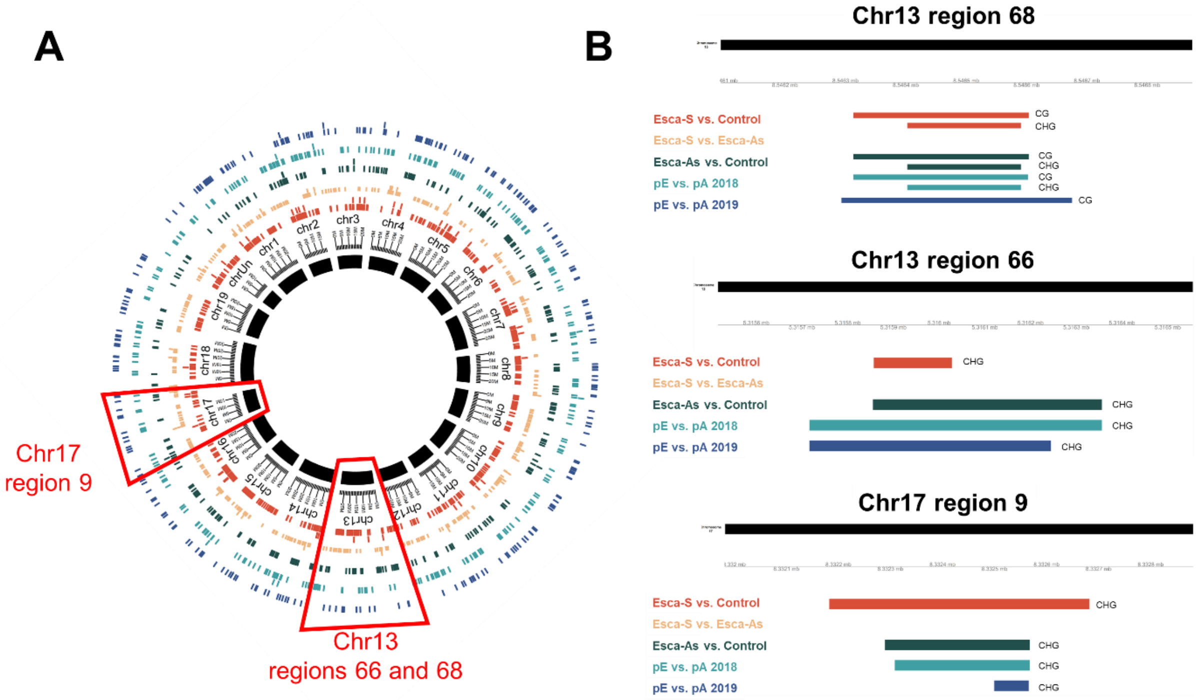

Figure S10. Graphical representation of the 'Enriched region' approach. (**A**) Circle plot representing the 19 grapevine chromosomes on which each line represent DMRs identified in all comparisons. From in to outer circle: ES-C (orange), ES-EA (yellow), EA-C (dark-green), pE-C 2018 (light blue)n pE-C 2019 (dark blue). (**B**) Representation of the overlapping DMRs identified at the three 'enriched' regions considered (see methods).

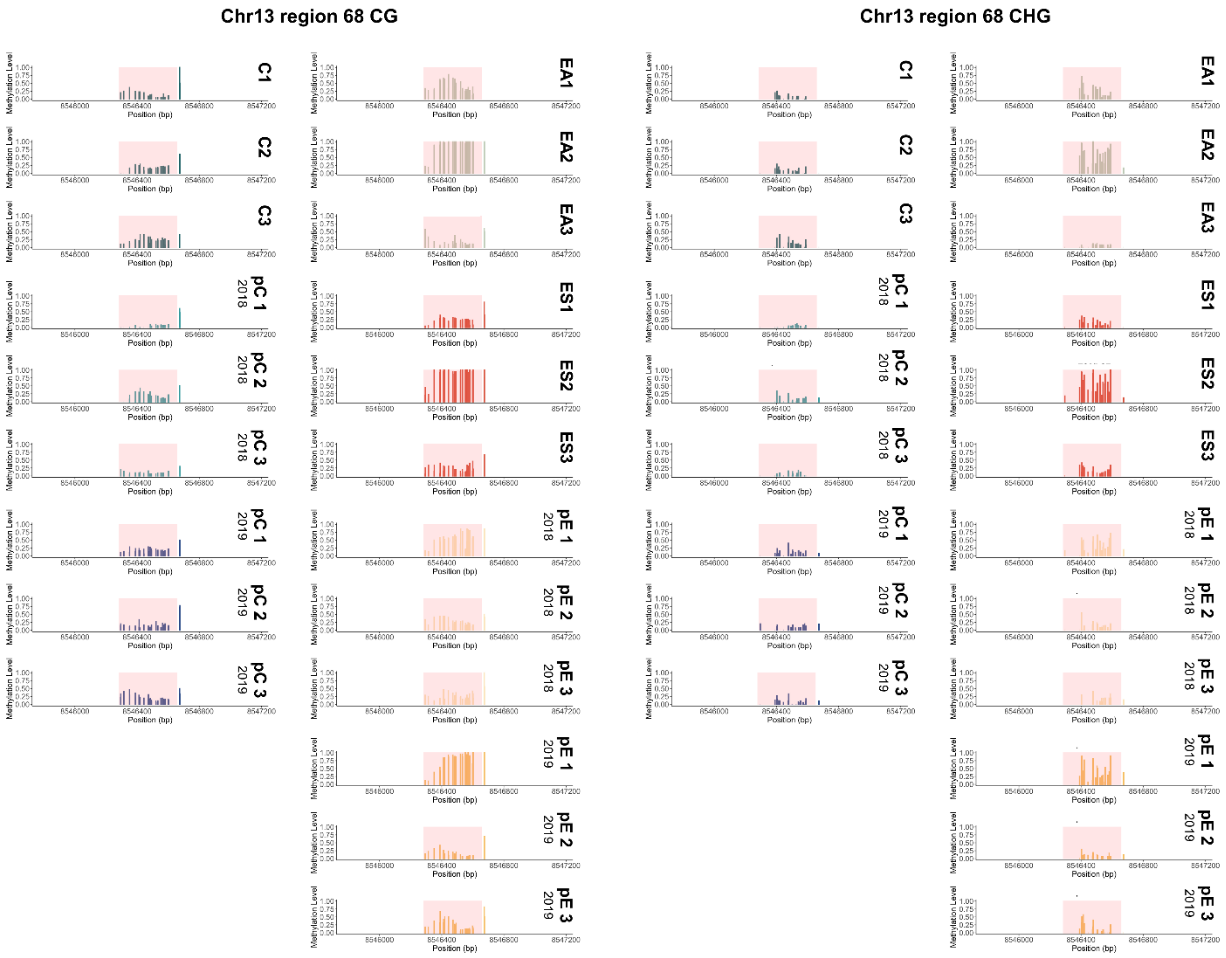

Figure S11. Details of the methylation distribution across region 68 in CG and CHG sequence context for each replicate. DMR position is highlighted by a semi-transparent red square.

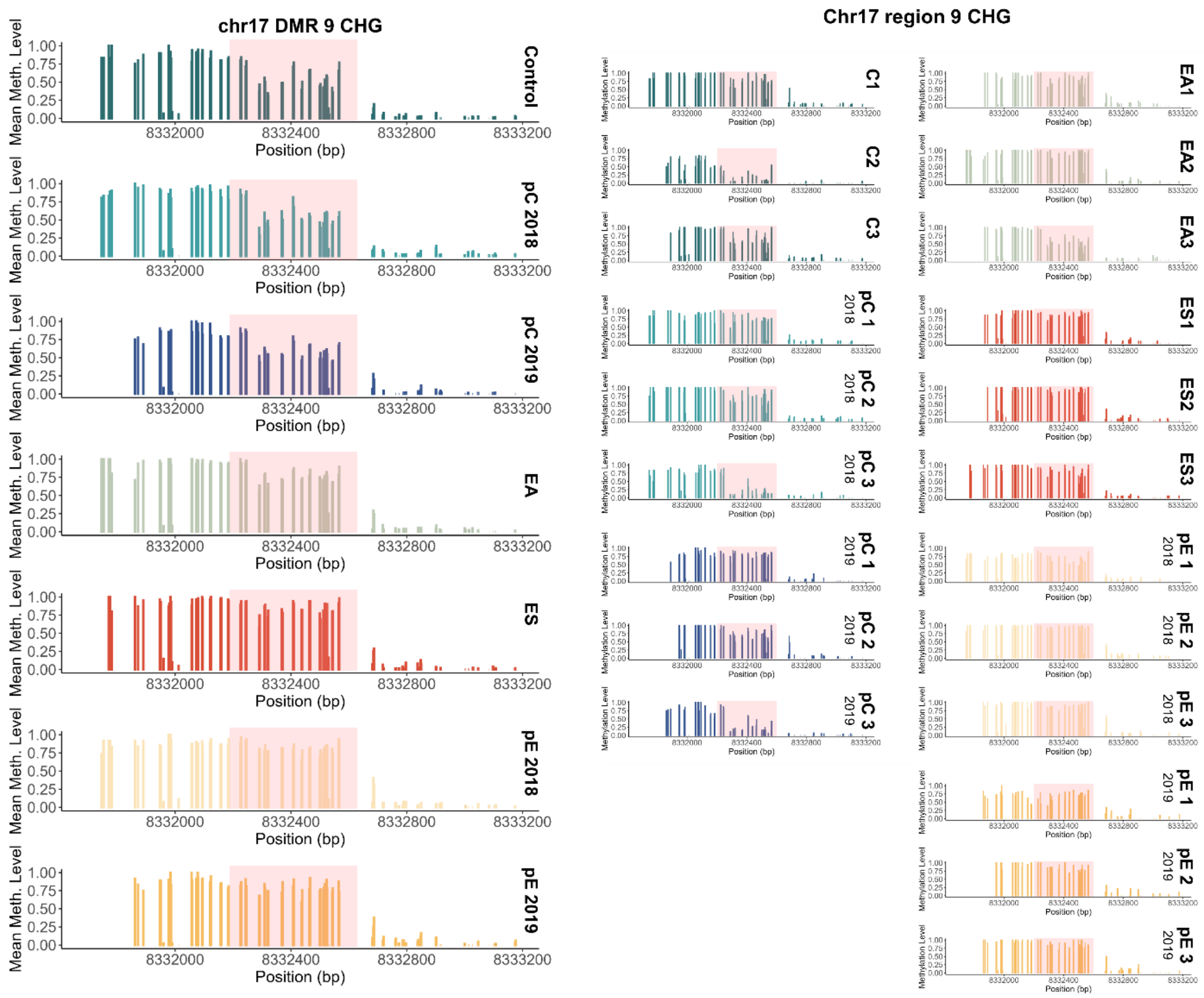

Figure S12. Methylation distribution across region 9 in the CHG sequence context. The left graphs show the mean methylation level at each cytosine position, calculated from the three replicates of each condition. The right graphs provide detailed methylation levels observed by sample. DMR positions are highlighted by a semi-transparent red square.

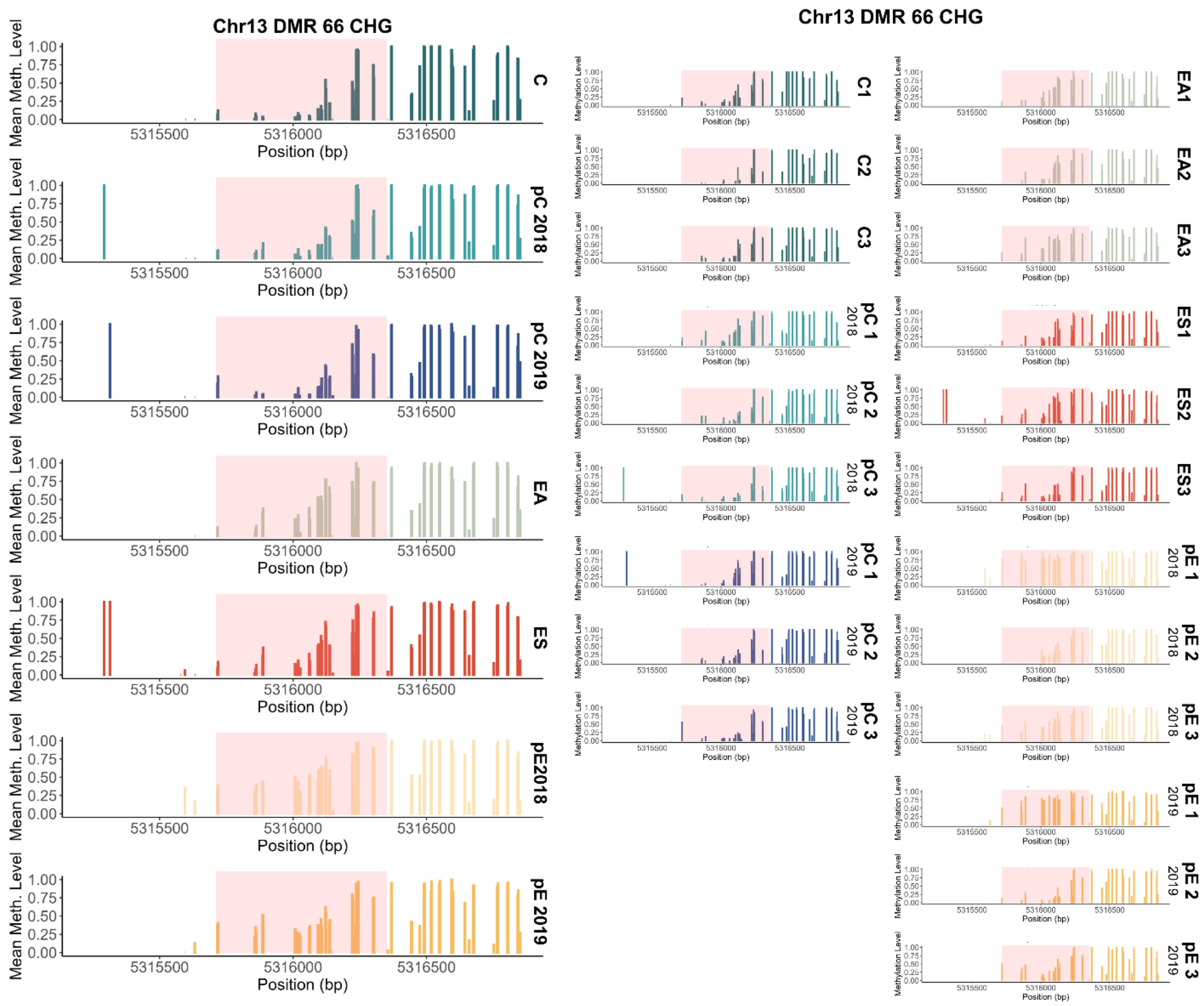

Figure S13. Methylation distribution across region 66 in the CHG sequence context. The left graph shows the mean methylation level at each cytosine position, calculated from the three replicates of each condition. The right graphs provide detailed methylation levels observed by sample. DMR positions are highlighted by a semi-transparent red square.

Table S1. Monitoring for Esca-symptoms expression from 2012 to 2019 for vineyard and greenhouse plants. Crosses indicate the symptomatic years, 'NA' indicate years for which no monitoring was performed, empty space means that the plant was asymptomatic during the concerned year. C: Control leaf, ES: esca-plant symptomatic leaf, EA: esca-plant asymptomatic leaf

| **Symptomatic years** | | | | | | | | |
|---|---|---|---|---|---|---|---|---|
| **Plant** | **2012** | **2013** | **2014** | **2015** | **2016** | **2017** | **2018** | **2019** |
| C 1 | | | | NA | NA | | | |
| C 2 | | | | NA | NA | | | |
| C 3 | | | | NA | NA | | | |
| ES 1 | | x | x | NA | NA | | x | x |
| ES 2 | | | x | NA | NA | x | | x |
| ES 3 | x | | | NA | NA | x | | x |
| pC 1 | | | | | | | | |
| pC 2 | | | | | | | | |
| pC 3 | | | | | | | | |
| pE 1 | x | x | x | | | x | x | x |
| pE 2 | x | x | x | | x | x | x | x |
| pE 3 | | x | x | | x | x | x | x |

Table S2. RNA-seq sequencing mapping and alignement results. C: Control leaf, ES: esca-plant symptomatic leaf, EA: esca-plant asymptomatic leaf

| Sample ID | Raw reads | M reads Mapped | % Mapped | % Successfully assigned alignements |
|---|---|---|---|---|
| C 1 | 4,1E+07 | 40.1 | 97.9 | 92.9 |
| C 2 | 4,1E+07 | 40.2 | 97.8 | 92.7 |
| C 3 | 4,1E+07 | 40.2 | 97.4 | 92.0 |
| EA 1 | 4,1E+07 | 39.7 | 96.4 | 92.5 |
| EA 2 | 4,1E+07 | 38.7 | 94.3 | 92.8 |
| EA 3 | 4,1E+07 | 31.2 | 76.1 | 91.7 |
| ES 1 | 4,1E+07 | 40.4 | 98.1 | 92.6 |
| ES 2 | 4,1E+07 | 39.8 | 97.4 | 92.3 |
| ES 3 | 4,1E+07 | 39.9 | 97.3 | 92.2 |

Table S3; Whole Genome Bisulfite Sequencing conversion rate, mapping results, and global methylation percentages observed in the three sequence context. C: Control leaf, ES: esca-plant symptomatic leaf, EA: esca-plant asymptomatic leaf, pC: Control leaf, pE: leaf sampled from esca-plant before symptom apparition.

| Sample | Plant status | Leaves symptom | % Bisulfite conversion | Raw reads | % Aligned | Mean cov | Total Cs | Total C's Methylated | % mCpG | % mCHG | % mCHH |
|---|---|---|---|---|---|---|---|---|---|---|---|
| C 1 | Control-plant | No | 99.2 | 89,969,322 | 58.3% | 21.1X | 1,685,448,493 | 209,012,004 | 59.0% | 29.8% | 3.9% |
| C 2 | Control-plant | No | 99.2 | 85,857,890 | 57.7% | 19.9X | 1,595,361,770 | 200,507,757 | 59.2% | 30.2% | 4.0% |
| C 3 | Control-plant | No | 99.2 | 85,874,048 | 57.3% | 19.8X | 1,597,434,035 | 204,329,560 | 58.0% | 30.3% | 4.2% |
| EA 1 | esca-plant | No | 99.2 | 88,744,070 | 58.0% | 21.2X | 1,682,489,848 | 218,278,488 | 59.7% | 30.8% | 4.0% |
| EA 2 | esca-plant | No | 99.2 | 85,693,142 | 59.8% | 21.7X | 1,673,190,005 | 210,464,427 | 59.2% | 30.0% | 4.0% |
| EA 3 | esca-plant | No | 99.2 | 88,002,221 | 59.1% | 21.4X | 1,682,095,309 | 212,669,987 | 58.8% | 30.2% | 4.1% |
| ES 1 | esca-plant | Yes | 99.3 | 86,919,650 | 60.8% | 20.7X | 1,685,068,619 | 204,197,905 | 59.3% | 29.6% | 3.9% |
| ES 2 | esca-plant | Yes | 99.2 | 86,799,936 | 61.8% | 20.6X | 1,726,257,469 | 217,832,477 | 60.5% | 30.3% | 4.1% |
| ES 3 | esca-plant | Yes | 99.2 | 87,995,267 | 60.4% | 20.9X | 1,703,221,662 | 216,224,047 | 61.9% | 31.0% | 4.0% |
| pC-2018 1 | Control-plant | No | 99.2 | 86,722,744 | 59.0% | 20.7X | 1,660,369,582 | 210,189,943 | 60.9% | 31.8% | 3.4% |
| pC-2018 2 | Control-plant | No | 99.1 | 84,833,118 | 58.8% | 20.1X | 1,637,594,842 | 209,133,931 | 61.0% | 31.7% | 3.4% |
| pC-2018 3 | Control-plant | No | 99.2 | 90,835,022 | 57.3% | 21.0X | 1,674,038,004 | 215,465,145 | 61.4% | 32.0% | 3.7% |
| pC-2019 1 | Control-plant | No | 99.2 | 86,209,061 | 58.1% | 20.3X | 1,649,230,044 | 210,192,700 | 62.0% | 33.1% | 2.8% |
| pC-2019 2 | Control-plant | No | 99.2 | 86,755,384 | 56.5% | 19.7X | 1,599,935,963 | 191,975,419 | 58.0% | 30.8% | 2.8% |
| pC-2019 3 | Control-plant | No | 99.1 | 88,865,482 | 58.1% | 21.0X | 1,697,587,098 | 203,780,553 | 59.4% | 30.7% | 2.7% |
| pE-2018 1 | esca-plant | No | 99.2 | 86,202,861 | 57.9% | 20.2X | 1,597,479,222 | 197,619,900 | 60.7% | 31.4% | 3.3% |
| pE-2018 2 | esca-plant | No | 99.1 | 86,643,785 | 59.0% | 20.7X | 1,678,948,922 | 225,631,236 | 63.2% | 33.4% | 3.9% |
| pE-2018 3 | esca-plant | No | 99.1 | 87,673,120 | 57.5% | 20.4X | 1,660,141,300 | 220,005,064 | 61.3% | 32.7% | 3.9% |
| pE-2019 1 | esca-plant | No | 99.2 | 87,911,229 | 58.4% | 20.7X | 1,646,565,882 | 180,896,013 | 57.3% | 28.7% | 2.4% |
| pE-2019 2 | esca-plant | No | 99.2 | 86,480,897 | 57.5% | 20.1X | 1,638,681,579 | 191,668,799 | 58.9% | 30.2% | 2.6% |
| pE-2019 3 | esca-plant | No | 99.2 | 87,656,807 | 57.3% | 20.4X | 1,658,485,342 | 188,730,949 | 57.0% | 29.1% | 2.5% |

Table S4. Average methylation observed for each condition across the three enriched regions identified. C: Control leaf, ES: esca-plant symptomatic leaf, EA: esca-plant asymptomatic leaf

| Region | Condition | Mean methylation level | sd | se | ci |
|---|---|---|---|---|---|
| | C | 0.23 | 0.31 | 0.03 | 0.06 |
| | 2018 pC | 0.25 | 0.31 | 0.03 | 0.06 |
| | 2019 pC | 0.24 | 0.3 | 0.03 | 0.06 |
| r66 | EA | 0.38 | 0.33 | 0.03 | 0.06 |
| | ES | 0.36 | 0.33 | 0.03 | 0.06 |
| | 2018 pE | 0.5 | 0.34 | 0.03 | 0.07 |
| | 2019 pE | 0.42 | 0.33 | 0.03 | 0.06 |
| | C | 0.14 | 0.13 | 0.01 | 0.02 |
| | 2018 pC | 0.07 | 0.1 | 0.01 | 0.02 |
| | 2019 pC | 0.14 | 0.12 | 0.01 | 0.02 |
| r68 | EA | 0.41 | 0.37 | 0.03 | 0.05 |
| | ES | 0.39 | 0.34 | 0.03 | 0.05 |
| | 2018 pE | 0.28 | 0.22 | 0.02 | 0.03 |
| | 2019 pE | 0.34 | 0.33 | 0.02 | 0.05 |
| | C | 0.53 | 0.35 | 0.03 | 0.07 |
| | 2018 pC | 0.54 | 0.33 | 0.03 | 0.06 |
| | 2019 pC | 0.57 | 0.32 | 0.03 | 0.06 |
| r9 | EA | 0.72 | 0.27 | 0.03 | 0.05 |
| | ES | 0.79 | 0.26 | 0.02 | 0.05 |
| | 2018 pE | 0.76 | 0.26 | 0.03 | 0.05 |
| | 2019 pE | 0.73 | 0.26 | 0.03 | 0.05 |

Table S5. Mean methylation levels observed for each region detailed per sample. C: Control leaf, ES: esca-plant symptomatic leaf, EA: esca-plant asymptomatic leaf

| region | Sample | mean_methylation | sd | se | ci |
|---|---|---|---|---|---|
| | C 1 | 0.27 | 0.31 | 0.05 | 0.1 |
| | C 2 | 0.14 | 0.29 | 0.05 | 0.1 |
| | C 3 | 0.26 | 0.31 | 0.05 | 0.11 |
| | EA 1 | 0.36 | 0.33 | 0.06 | 0.11 |
| | EA 2 | 0.35 | 0.34 | 0.06 | 0.12 |
| | EA 3 | 0.43 | 0.31 | 0.05 | 0.11 |
| | ES 1 | 0.31 | 0.31 | 0.05 | 0.11 |
| | ES 2 | 0.44 | 0.34 | 0.06 | 0.11 |
| | ES 3 | 0.32 | 0.32 | 0.05 | 0.11 |
| | 2018_pC1 | 0.34 | 0.32 | 0.05 | 0.11 |
| r66 | 2019_pC1 | 0.29 | 0.35 | 0.06 | 0.12 |
| | 2018_pC2 | 0.22 | 0.3 | 0.05 | 0.1 |
| | 2019_pC2 | 0.22 | 0.29 | 0.05 | 0.1 |
| | 2018_pC3 | 0.18 | 0.28 | 0.05 | 0.1 |
| | 2019_pC3 | 0.21 | 0.27 | 0.05 | 0.09 |
| | 2018_pE1 | 0.74 | 0.23 | 0.04 | 0.08 |
| | 2019_pE1 | 0.66 | 0.23 | 0.04 | 0.08 |
| | 2018_pE2 | 0.3 | 0.32 | 0.05 | 0.11 |
| | 2019_pE2 | 0.23 | 0.31 | 0.05 | 0.1 |
| | 2018_pE3 | 0.45 | 0.31 | 0.05 | 0.11 |
| | 2019_pE3 | 0.37 | 0.3 | 0.05 | 0.1 |
| | C 1 | 0.12 | 0.16 | 0.02 | 0.04 |
| | C 2 | 0.13 | 0.12 | 0.02 | 0.03 |
| | C 3 | 0.17 | 0.12 | 0.02 | 0.03 |
| | EA 1 | 0.34 | 0.19 | 0.03 | 0.05 |
| | EA 2 | 0.8 | 0.3 | 0.04 | 0.08 |
| | EA 3 | 0.1 | 0.13 | 0.02 | 0.03 |
| | ES 1 | 0.19 | 0.14 | 0.02 | 0.03 |
| | ES 2 | 0.78 | 0.28 | 0.04 | 0.07 |
| | ES 3 | 0.19 | 0.13 | 0.02 | 0.03 |
| | 2018_pC1 | 0.04 | 0.1 | 0.01 | 0.03 |
| r68 | 2019_pC1 | 0.15 | 0.11 | 0.01 | 0.03 |
| | 2018_pC2 | 0.12 | 0.12 | 0.02 | 0.03 |
| | 2019_pC2 | 0.12 | 0.14 | 0.02 | 0.04 |
| | 2018_pC3 | 0.05 | 0.07 | 0.01 | 0.02 |
| | 2019_pC3 | 0.16 | 0.11 | 0.01 | 0.03 |
| | 2018_pE1 | 0.44 | 0.22 | 0.03 | 0.06 |
| | 2019_pE1 | 0.66 | 0.31 | 0.04 | 0.08 |
| | 2018_pE2 | 0.17 | 0.14 | 0.02 | 0.04 |
| | 2019_pE2 | 0.12 | 0.14 | 0.02 | 0.04 |
| | 2018_pE3 | 0.22 | 0.17 | 0.02 | 0.05 |
| | 2019_pE3 | 0.21 | 0.18 | 0.02 | 0.05 |

| | | | | | |
|---|---|---|---|---|---|
| | C 1 | 0.76 | 0.25 | 0.04 | 0.09 |
| | C 2 | 0.17 | 0.19 | 0.03 | 0.06 |
| | C 3 | 0.65 | 0.26 | 0.04 | 0.09 |
| | EA 1 | 0.76 | 0.25 | 0.04 | 0.09 |
| | EA 2 | 0.85 | 0.25 | 0.04 | 0.09 |
| | EA 3 | 0.56 | 0.24 | 0.04 | 0.08 |
| | ES 1 | 0.79 | 0.23 | 0.04 | 0.08 |
| | ES 2 | 0.86 | 0.26 | 0.04 | 0.09 |
| | ES 3 | 0.71 | 0.26 | 0.04 | 0.09 |
| | 2018_pC1 | 0.67 | 0.24 | 0.04 | 0.08 |
| r9 | 2019_pC1 | 0.73 | 0.23 | 0.04 | 0.08 |
| | 2018_pC2 | 0.69 | 0.25 | 0.04 | 0.09 |
| | 2019_pC2 | 0.71 | 0.24 | 0.04 | 0.08 |
| | 2018_pC3 | 0.25 | 0.28 | 0.05 | 0.1 |
| | 2019_pC3 | 0.28 | 0.27 | 0.05 | 0.09 |
| | 2018_pE1 | 0.6 | 0.21 | 0.04 | 0.07 |
| | 2019_pE1 | 0.6 | 0.22 | 0.04 | 0.08 |
| | 2018_pE2 | 0.86 | 0.26 | 0.04 | 0.09 |
| | 2019_pE2 | 0.77 | 0.25 | 0.04 | 0.09 |
| | 2018_pE3 | 0.81 | 0.24 | 0.04 | 0.08 |
| | 2019_pE3 | 0.8 | 0.27 | 0.05 | 0.09 |
